# Microglia-astrocyte crosstalk regulates synapse remodeling via Wnt signaling

**DOI:** 10.1101/2024.02.08.579178

**Authors:** Travis E. Faust, Yi-Han Lee, Ciara D. O’Connor, Margaret A. Boyle, Georgia Gunner, Ana Badimon, Pinar Ayata, Anne Schaefer, Dorothy P. Schafer

## Abstract

Astrocytes and microglia are emerging key regulators of activity-dependent synapse remodeling that engulf and remove synapses in response to changes in neural activity. Yet, the degree to which these cells communicate to coordinate this process remains an open question. Here, we use whisker removal in postnatal mice to induce activity-dependent synapse removal in the barrel cortex. We show that astrocytes do not engulf synapses in this paradigm. Instead, astrocytes reduce their contact with synapses prior to microglia-mediated synapse engulfment. We further show that reduced astrocyte-contact with synapses is dependent on microglial CX3CL1-CX3CR1 signaling and release of Wnts from microglia following whisker removal. These results demonstrate an activity-dependent mechanism by which microglia instruct astrocyte-synapse interactions, which then provides a permissive environment for microglia to remove synapses. We further show that this mechanism is critical to remodel synapses in a changing sensory environment and this signaling is upregulated in several disease contexts.

## Introduction

Microglia and astrocytes are now appreciated as key regulators of activity-dependent synapse remodeling in the developing brain. In the developing retinogeniculate circuit, microglia and astrocytes engulf and remove synapses from less active neurons.^1–3^ In the cortex, spinal cord, and thalamus, IL-33 from astrocytes has been shown to stimulate microglial synaptic pruning.^4–6^ Still, it remains unknown if neural activity can regulate this glial cell cross talk to regulate synapse remodeling and it is unknown if and how microglia signal to astrocytes to coordinate this process.

The somatosensory system is widely used to study activity-dependent synaptic remodeling.^7, 8^ In humans, loss of sensory input (e.g. amputation) leads to a reduction in the cortical representation of the affected region.^9^ Similarly, in rodents, removal of sensory input (e.g. vision, hearing, somatosensation) leads to remodeling of this representation in the CNS. This includes trimming, plucking or cauterizing whiskers, which can lead to remodeling of synaptic connections in a region of the somatosensory cortex called the barrel cortex. In the rodent barrel cortex, it is known that removal of whiskers during a critical window in postnatal development (postnatal day 1-3; P1-P3) results in a failure of thalamocortical synapses to form an appropriate synaptic representation of the whiskers on the snout.^10–12^ We have since shown that after this critical window, whisker removal by cauterization or trimming in postnatal day 4 (P4) mice elicits removal of previously formed layer IV thalamocortical (TC) synapses by phagocytic microglia.^13^ Unlike in the visual thalamus,^3^ this engulfment of presynaptic terminals by microglia was not dependent on complement receptor 3. Instead, microglia-mediated synapse removal was mediated by neuronal fractalkine (CX3CL1)-microglial fractalkine receptor (CX3CR1) signaling. In mice deficient in CX3CL1 or CX3CR1, whisker lesioning by cauterization in P4 mice no longer induced TC synapse elimination in the corresponding barrel cortex. Earlier work demonstrated that deficient CX3CR1 signaling in microglia affects microglia-mediated synapse pruning and maturation,^14–17^ however CX3CR1 is a G-protein coupled receptor (GPCR), not an engulfment receptor, and the mechanism by which this receptor regulates microglia function at synapses remained unknown.

Here, we find that whisker lesioning in postnatal mice, which induces microglia to engulf and remove synapses, does not induce astrocyte engulfment of TC synapses. Instead, astrocyte processes decrease their physical association with synapses as microglia begin to engulf and remove synapses. We then identify that this reduction in astrocyte process coverage of synapses is dependent on microglial CX3CL1-CX3CR1 signaling upstream of Wnt release from microglia and activation of Wnt receptor signaling in astrocytes. Genetic ablation of CX3CL1-CX3CR1 signaling or microglial Wnt release results in a failure of astrocytes to upregulate Wnt receptor signaling and reduce their contact with TC synapses following whisker lesioning, as well as inhibition of TC synapse removal by microglia. We further identify that transcriptional activation of Wnt receptor signaling in astrocytes is common across multiple neuroinflammatory and neurodegenerative conditions where synapse loss is a hallmark feature. These results identify a mechanism by which microglia and astrocytes communicate to coordinate synapse remodeling in response to changes in neural activity in the developing brain. In the process, we also identify a previously unidentified mechanism by which CX3CL1-CX3CR1 signaling regulates microglia-mediated synapse remodeling via Wnt release, which has relevance to multiple neurological diseases.

## Results

### Astrocytes decrease their association with thalamocortical synapses following whisker lesioning

Our previous work showed that microglia engulf thalamocortical (TC) presynaptic inputs in the postnatal barrel cortex in response to whisker lesioning.^13^ As astrocytes have also been shown to engulf synapses in other brain regions,^7^ we first assessed whether astrocytes were similarly engulfing these TC inputs in this paradigm. All whiskers were lesioned by cauterization on one side of the P4 mouse snout such that each animal had its own internal control (intact whiskers on the other side of the snout) (Figure 1A). Importantly, P4 is after the closure of the more stereotypical barrel cortex critical window in which whisker removal prevents the normal initial wiring of the somatotopic synaptic map.^10–12^ In agreement with our previous work,^13^ within 6 days of P4 whisker lesioning, vesicular glutamate transporter 2 (VGluT2)^+^ TC presynaptic terminals within layer IV of the barrel cortex were reduced (Figure 1B-C). There was also a concomitant increase in engulfed VGluT2^+^ material within microglial lysosomes within 5 days of whisker lesioning (Figure S1A-B). In contrast, there was no significant increase in engulfed VGluT2^+^ material within astrocytes lysosomes (Figure 1D-F).

**Figure 1.**
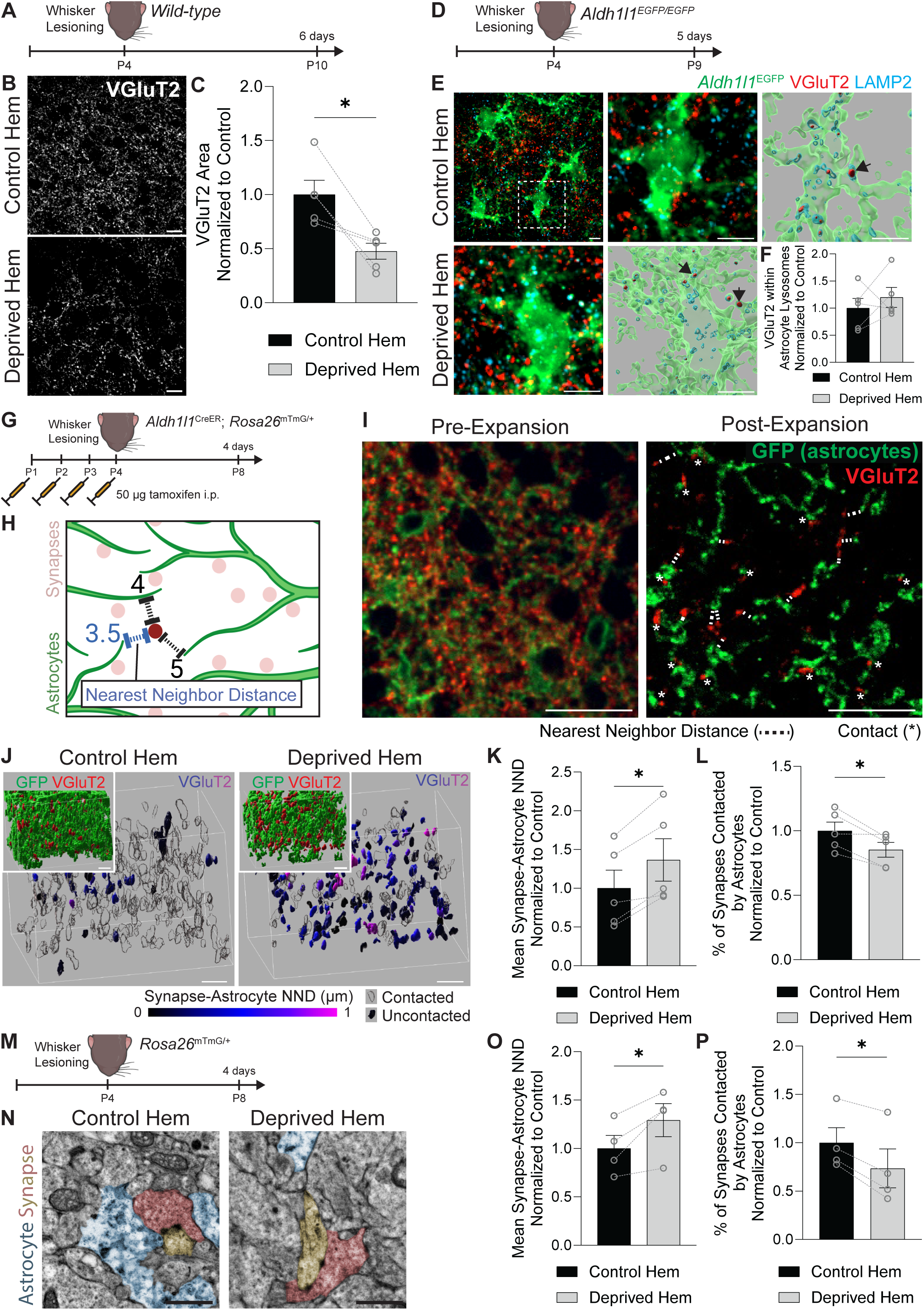
Astrocytes decrease their association with thalamocortical synapses following whisker lesioning. (A) Diagram of experimental protocol used to assess remodeling of thalamocortical synaptic terminals in wild-type mice at 6 days after unilateral whisker lesioning. (B) Representative fluorescent images of anti-VGluT2^+^ thalamocortical synaptic terminals in layer IV of the barrel cortex in the control and deprived hemispheres at 6 days post-lesioning. Scale bars 10 µm. (C) Quantification of the density of anti-VGluT2^+^ thalamocortical synaptic terminals in layer IV of the barrel cortex at 6 days post-lesioning shows reduced density in the deprived hemisphere (hem) compared to the control hem (Ratio paired t test: *n* = 5 mice. * *p* < 0.05). (D) Diagram of experimental protocol used to assess astrocyte engulfment of thalamocortical synaptic terminals in *Aldh1l1^EGFP/EGFP^* mice at 5 days after unilateral whisker lesioning. (E) Representative max intensity projections of anti-VGluT2^+^ thalamocortical synaptic terminals (red), anti-LAMP2^+^ lysosomes (cyan), and *Aldh1l1^EGFP/EGFP^* astrocytes immunolabeled with anti-GFP (green) in layer IV of the barrel cortex in control (top left; inset top middle) and deprived (bottom left) hems at 5 days post-lesioning. Representative Imaris surface reconstructions of anti-VGluT2^+^ synaptic material (red) contained within anti-LAMP2^+^ lysosomes (cyan) inside *Aldh1l1^EGFP/EGFP^* astrocytes immunolabeled with anti-GFP (green) in control (top right) and deprived (bottom middle) hems. Arrows indicate examples of VGluT2^+^ synaptic material contained within LAMP2^+^ astrocyte lysosomes. Scale bars 5 µm. (F) Quantification of anti-VGluT2^+^ synaptic material within astrocyte lysosomes at 5 days after whisker lesioning does not show a difference between the deprived hem and the control hem of the barrel cortex (Ratio paired t test: *n* = 5 mice). (G) Diagram of experimental protocol used for expansion microscopy analysis of *Aldh1l1^CreER/+^*_;_ *Rosa26^mTmG/+^* mice at 4 days after unilateral whisker lesioning. (H) Diagram of the distances between a given synapse (dark red) and the nearest astrocyte processes (green), the shortest of which (blue) is considered the nearest neighbor distance. (I) Representative immunofluorescent images of layer IV barrel cortex brain sections from a *Aldh1l1^CreER/+^*_;_ *Rosa26^mTmG/+^* mouse, immunolabeled with anti-VGluT2 to label thalamocortical synapses (red) and anti-GFP to label astrocyte membranes (green). The left image shows the tissue before gel-embedding for expansion microscopy (pre-expansion) and the right image shows the tissue after ∼4-fold expansion for expansion microscopy. Asterisks (*) indicate synapses contacted by astrocytes. For synapses without direct contact, a dashed line (---) is drawn between the synapse and the nearest astrocyte to indicate the nearest neighbor distance (NND). Scale bars 20 µm (not adjusted by expansion index). (J) Representative Imaris surface reconstructions of expansion microscopy z-stack images from layer IV barrel cortex from the control and deprived hemispheres of *Aldh1l1^CreER/+^*_;_ *Rosa26^mTmG/+^* mice at 4 days after unilateral whisker lesioning. Anti-VGluT2^+^ thalamocortical synaptic terminals are pseudo-colored by the nearest neighbor distance (NND; corrected for expansion index) to the nearest astrocyte process. Anti-VGluT2^+^ surfaces with NND of 0 are considered “contacted” and are shown in transparent black. Insets show anti-GFP^+^ astrocytes (green) and anti-VGluT2^+^ thalamocortical synaptic terminals (red). Scale bars 2 µm (corrected for expansion index). (K) Quantification of the mean synapse-astrocyte NND in expansion microscopy images of layer IV of the barrel cortex of *Aldh1l1^CreER/+^*_;_ *Rosa26^mTmG/+^* mice at 4 days after unilateral whisker lesioning shows increased NND in the deprived hem compared to the control hem (Ratio paired t test: n = 5 mice. * p < 0.05). (L) Quantification of the percentage of synapses contacted by astrocytes in expansion microscopy images of layer IV of the barrel cortex of *Aldh1l1^CreER/+^*_;_ *Rosa26^mTmG/+^* mice at 4 days after unilateral whisker lesioning shows a reduced contact percentage in the deprived hem compared to the control hem (Ratio paired t test: n = 5 mice. * p < 0.05). (M) Diagram of experimental protocol used for electron microscopy analysis of *Rosa26^mTmG/+^* mice at 4 days after unilateral whisker lesioning. (N) Representative electron microscopy images of synapses (pseudocolored red and yellow) and astrocytes (pseudocolored blue) in the control and deprived hems. Scale bars 1 µm. (O) Quantification of the mean synapse-astrocyte NND in electron microscopy images of layer IV of the barrel cortex at 4 days after unilateral whisker lesioning shows increased NND in the deprived hem compared to the control hem (Ratio paired t test: n = 4 mice. * p < 0.05). (P) Quantification of the percentage of synapses contacted by astrocytes in electron microscopy images of layer IV of the barrel cortex at 4 days after unilateral whisker lesioning shows a reduced contact percentage in the deprived hem compared to the control hem (Ratio paired t test: n = 4 mice. * p < 0.05). All data are presented as mean ± SEM. See also Figure S1.

Astrocytes are known to have an extensive network of fine processes which closely associate with neuronal synapses. We therefore assessed whether astrocyte process proximity to synapses, versus engulfment, could instead be impacted by whisker lesioning. To assess the apposition of astrocyte processes to synapses in greater detail, we generated transgenic mice in which astrocytes were labeled with membrane-bound GFP (mGFP) (*Aldh1l1^CreER/+^*; *Rosa26^mTmG/+^*) (Figure S1C-E).^18, 19^ To enhance the imaging resolution of these mGFP^+^ fine astrocytic processes, we used expansion microscopy (Figure S1F). On average, we achieved a 4-fold expansion of the tissue (Figure S1G), thereby providing a 4-fold increase in resolution. We first assessed the degree of astrocyte process contact with synapses in the barrel cortex of *Aldh1l1^CreER/+^*; *Rosa26^mTmG/+^* mice at 4 days after whisker lesioning (lesioning at P4 and assessment at P8) in expanded tissue (Figure 1G-I). Compared to the control hemisphere, the average distance between a VGluT2^+^ TC presynaptic terminal and its nearest astrocyte process (nearest neighbor distance, NND) was increased (Figure 1H-K) and the percentage of VGluT2^+^ synapses contacted by astrocytes was decreased in the deprived hemisphere (Figure 1L). We further confirmed these findings using electron microscopy (Figure 1M-P). This was concomitant with a reduction in astrocyte process density (Figure S1H-I) with no change in total astrocyte numbers in layer IV of the barrel cortex (Figure S1J-K). These data demonstrate that astrocytes in the barrel cortex do not engulf synapses but rather respond to whisker lesioning by reducing their physical interaction with synapses just prior to microglia-mediated synapse removal.

### Decreased association of astrocyte processes with thalamocortical synapses is CX3CL1-dependent

We next sought to test whether decreased astrocyte-synapse interactions following whisker lesioning required signaling from microglia. We previously found that neuronal CX3CL1 signaling to microglial CX3CR1 was critical for microglia to engulf and remove synapses.^13^ CX3CL1 is the only known *in vivo* ligand of CX3CR1, which is a receptor exclusively expressed by microglia and other myeloid-derived cells in the brain at P4. However, CX3CR1 is not an engulfment receptor. Instead, it is a G-protein coupled chemokine receptor. We previously showed that loss of CX3CR1 or CX3CL1 had no effect on the number of microglia in the barrel cortex following whisker lesioning, but it did block microglia-mediated engulfment and removal of TC synapses.^13^ We reasoned that there could be signaling downstream of CX3CL1-CX3CR1 that could modulate communication between microglia and astrocytes to influence synapse remodeling (Figure 2A). Because CX3CR1-deficient mice (*Cx3cr1^-/-^*; also referred to as *Cx3cr1^EGFP/EGFP^*)^20^ are engineered to express GFP in microglia, which would overlap with mGFP signal in astrocytes in *Aldh1l1^CreER/+^; Rosa26^mTmG/+^* mice, we tested this possibility in mice deficient in the ligand CX3CL1 (*Cx3cl1^-/-^*).^21^ Consistent with CX3CL1-CX3CR1 signaling in microglia being upstream of changes in astrocyte-synapse interactions following whisker lesioning, we first found astrocytes no longer decreased their process density in response to P4 whisker lesioning in CX3CL1-deficient mice (*Cx3cl1^-/-^; Aldh1l1^CreER/+^*; *Rosa26^mTmG/+^* mice) (Figure 2B-C). Similarly, astrocytes no longer decreased their association with TC synapses in the barrel cortex in CX3CL1-deficient mice (Figure 2D-F). As CX3CL1 is highly selective for CX3CR1 on microglia, these experiments suggest that decreased astrocyte-synapse association after whisker lesioning is due to CX3CL1-CX3CR1-dependent microglial signaling prior to synapse engulfment. They further suggest that there is a CX3CL1-CX3CR1-dependent signal from microglia that elicits astrocyte process remodeling following whisker lesioning in neonates.

**Figure 2:**
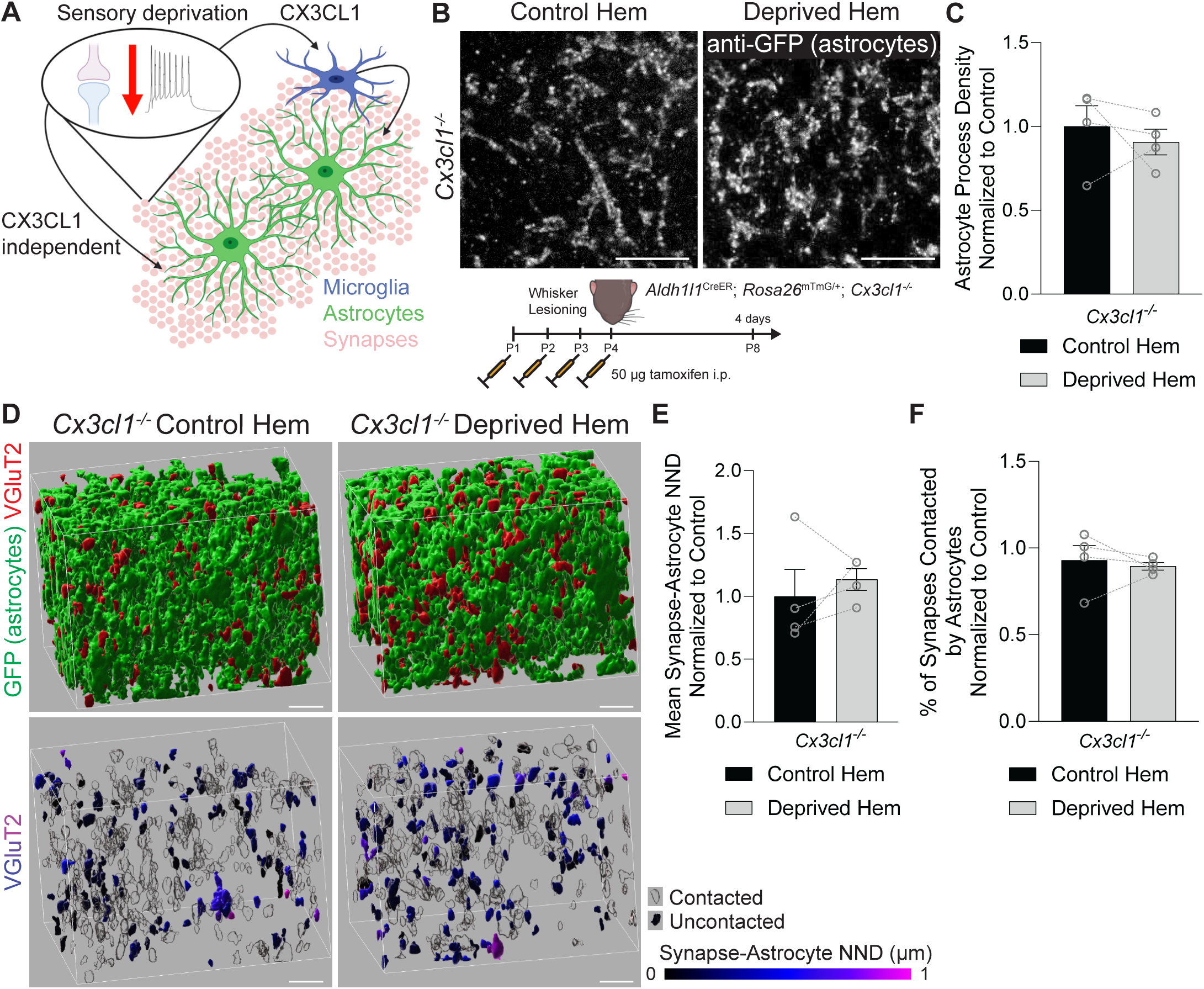
Decreased association of astrocyte processes with thalamocortical synapses is CX3CL1-dependent. (A) Schematic of potential astrocyte signaling mechanisms after sensory deprivation: CX3CL1-dependent microglia-astrocyte signaling and CX3CL1-independent neuron-astrocyte signaling. (B) Top: Representative fluorescent images of expanded layer IV barrel cortex brain sections from *Aldh1l1^CreER/+^*_;_ *Rosa26^mTmG/+^; Cx3cl1^-/-^* mice collected 4 days after unilateral whisker lesioning. Sections from the control and deprived hemispheres are labeled with anti-GFP to identify astrocyte processes. Scale bars 5 µm (corrected for expansion index). Bottom: diagram of experimental protocol used for expansion microscopy analysis of *Aldh1l1^CreER/+^*_;_ *Rosa26^mTmG/+^; Cx3cl1^-/-^*mice at 4 days after unilateral whisker lesioning. (C) Quantification of astrocyte process density in layer IV of the barrel cortex at 4 days after whisker lesioning does not show differences between the deprived and control hemispheres (hem) (Ratio paired t test: n = 4 mice). (D) Representative Imaris surface reconstructions of anti-VGluT2^+^ thalamocortical synaptic terminals in layer IV barrel cortex from the control and deprived hemispheres of *Aldh1l1^CreER/+^*_;_ *Rosa26^mTmG/+^; Cx3cl1^-/-^*mice at 4 days after unilateral whisker lesioning. Top: anti-GFP^+^ astrocytes (green) and anti-VGluT2^+^ thalamocortical synaptic terminals (red). Bottom: anti-VGluT2^+^ thalamocortical synaptic terminals pseudo-colored by the nearest neighbor distance (NND; corrected for expansion index) to the nearest astrocyte process. Anti-VGluT2^+^ surfaces with NND of 0 are considered “contacted” and are shown in transparent black. Scale bars 2 µm (corrected for expansion index). (E) Quantification of the mean synapse-astrocyte NND in layer IV of the barrel cortex of *Aldh1l1^CreER/+^*_;_ *Rosa26^mTmG/+^; Cx3cl1^-/-^* mice at 4 days after unilateral whisker lesioning does not show a difference in NND between deprived and control hems (Ratio paired t test: n = 4 mice). (F) Quantification of the percentage of synapses contacted by astrocytes in layer IV of the barrel cortex of *Aldh1l1^CreER/+^*_;_ *Rosa26^mTmG/+^; Cx3cl1^-/-^*mice at 4 days after unilateral whisker lesioning does not show a difference in the contact percentage between the deprived and control hems (Ratio paired t test: *n* = 4 mice). All data are presented as mean ± SEM.

### Whisker lesioning induces Wnt receptor signaling in astrocytes

To interrogate putative CX3CL1-CX3CR1-dependent microglia-to-astrocyte signaling following whisker lesioning, we performed cell type-specific translating ribosome affinity purification followed by bulk RNA sequencing (TRAP-Seq). For these experiments, astrocyte ribosomes were labeled using transgenic *Aldh1l1^EGFP-L10a/+^* mice^22^ (Figure 3A) and microglial ribosomes were labeled using *Cx3cr1^CreER/+ (Litt)^; Eef1a1^LSL-EGFP-L10a/+^* mice^23^ (Figure 3B). Then, 24 hours after unilateral whisker lesioning in P4 mice, the deprived and non-deprived barrel cortices were micro-dissected and GFP-tagged ribosomes were immunoprecipitated with a GFP antibody followed by RNA-seq. As a quality control, we insured that cell-type specific genes for microglia and astrocytes were highly enriched in the TRAP-bound RNA versus the unbound fraction and we confirmed that genes for other brain cell types were not enriched (Figure S2). Comparing the deprived and non-deprived cortices, we identified 147 genes that were upregulated and 39 genes that were downregulated in astrocyte TRAP-bound RNA in the deprived barrel cortex (Figure 3C), as well as 471 genes that were upregulated and 156 genes that were downregulated in microglia TRAP-bound RNA in the deprived barrel cortex (Figure 3D). To identify potential signaling pathways upstream of the astrocyte transcriptional changes, we performed Ingenuity Pathway Analysis (IPA) of higher order root regulators (Figure 3E) and direct upstream regulators (Figure 3F). IPA identified many regulators related to the canonical Wnt signaling pathway including β-catenin (CTNNB1) and its co-activated transcription factors T-cell factor/lymphoid enhancer factor (TCF/LEF). We then used NicheNet^24^ to perform ligand-receptor analysis between microglia and astrocytes (Figure 3G). For this analysis, we took all ligand-receptor pairs that are expressed in microglia and astrocytes independent of whisker lesioning and probed which microglia ligands have the strongest regulatory association with the genes that are transcriptionally increased or decreased in astrocytes following whisker lesioning. Similar to IPA, several microglia-derived Wnt ligands were identified among the top putative ligand-receptor interactions (Figure 3H). These data suggest that microglia-released Wnts may induce transcriptional changes observed in astrocytes after whisker lesioning. Interestingly, when we compared our astrocyte sequencing results to published astrocyte TRAP-Seq datasets from other conditions in which changes in astrocyte association with synapses, microglial synapse engulfment, and decreases in synapses have been documented (day vs. night, aging, Alzheimer’s disease, etc.)^25–29^ we found similar evidence of astrocyte Wnt receptor signaling by NicheNet (Figure 3I-J). These data support that Wnt signaling in astrocytes could be a more generalized mechanism involved in regulating microglia-astrocyte crosstalk and synapse remodeling.

**Figure 3.**
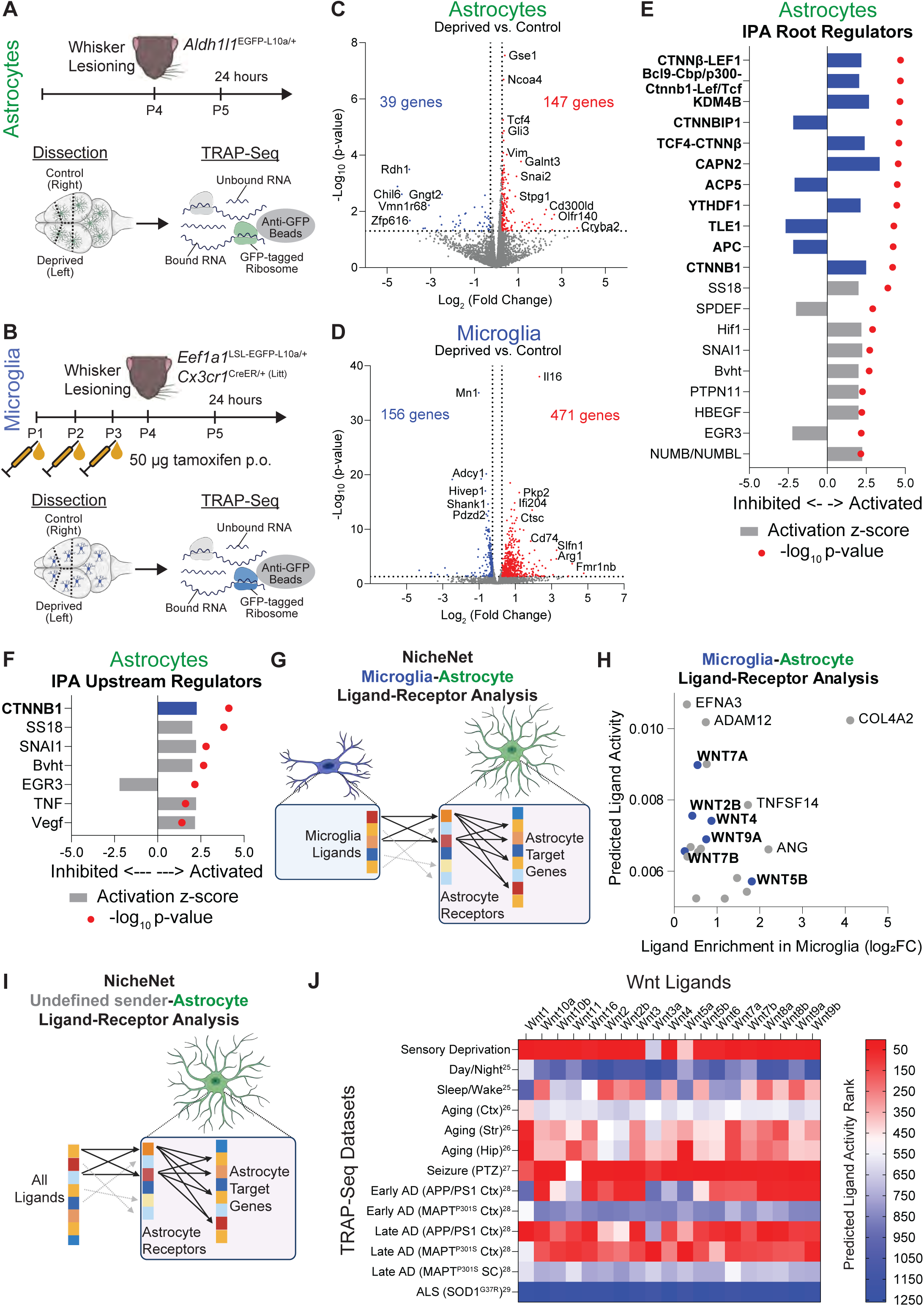
Whisker lesioning induces Wnt signaling in astrocytes. (A-B) Diagram of experimental protocol used for TRAP-Seq analysis of ribosome-bound RNA from (A) astrocytes and (B) microglia isolated from the control and deprived hemispheres (hem) of the primary somatosensory cortex at 24 hours after unilateral whisker lesioning. (C-D) Volcano plots of changes in gene expression in (C) astrocytes and (D) microglia in the deprived hem vs. the control hem of the barrel cortex at 24 hours after whisker lesioning. Plots include all genes with significant enrichment in the TRAP-enriched fraction vs. the unbound fraction. Genes with |fold-change| > 1.2 and p-value < 0.05 are considered differentially expressed genes (DEGs) for downstream analyses. DEGs increased in the deprived hem are colored red and DEGs decreased in the deprived hem are colored blue. (E-F) Ingenuity pathway analysis (IPA) of (E) root regulators and (F) upstream regulators for DEGs in deprived hem astrocytes. Bars indicate the activation z-score and red dots indicate the p-value of activation score. Root and upstream regulators involved in the Wnt signaling pathway are bolded and highlighted in blue. (G) Graphical overview of NicheNet ligand-receptor analysis of microglia signaling to astrocytes after whisker lesioning. TRAP-Seq data was used for the analysis, selecting potential ligands from the list of microglia-enriched genes, potential receptors from the list of astrocyte-enriched genes, and target genes from the list of astrocyte DEGs. (H) Plot of gene enrichment in microglia vs. the NicheNet predicted ligand activity for the top 20 ligands predicted by NicheNet. Wnt ligands are bolded and highlighted in blue. (I) Graphical overview of NicheNet ligand-receptor analysis of public astrocyte TRAP-Seq datasets. For the analysis, all ligands in the NicheNet database were used (not restricted by cell-type), receptors were selected from the list of astrocyte-enriched genes within each dataset, and astrocyte DEGs were used as the target genes. (J) Heatmap showing the rank of the predicted ligand activity of Wnt ligands in different astrocyte TRAP-Seq datasets. Superscripts indicate the reference numbers listed in the main text. See also Figure S2.

### Increased Wnt receptor signaling in layer IV astrocytes is CX3CL1/CX3CR1-dependent

Using the barrel cortex whisker lesioning paradigm, we next sought to validate our findings of Wnt receptor activation in astrocytes. Two of the major Wnt signaling-related root regulators in astrocytes identified by IPA were β-catenin and TCF/LEF. Upon binding of Wnt ligand to its receptors on the plasma membrane, β-catenin translocates to the nucleus and subsequently induces TCF/LEF-mediated transcriptional regulation of Wnt-responsive genes (Figure 4A). To test whether whisker lesioning induces TCF/LEF activity in astrocytes, we used the *TCF/Lef:H2B-GFP* reporter mouse line in which activation of a multimerized TCF/LEF response element drives expression of nuclear GFP (Figure 4B).^30^ Using immunofluorescence confocal microscopy, we confirmed canonical Wnt signaling induction in layer IV astrocytes (identified by SOX9^+^ nuclei) at 24 hours following P4 whisker lesioning as indicated by a significant increase in *TCF/Lef:H2B-GFP* reporter intensity in the deprived barrel cortex (Figure 4C-D). Interestingly, we also observed *TCF/Lef:H2B-GFP* signal in non-astrocytes. However this signal may have been due to the use of an artificial multimerized TCF reporter line.^31^ We, therefore, also performed immunostaining for β-catenin and analyzed levels of β-catenin in nuclei, similar to previous studies^32^ (Figure 4E). In agreement with our results from *TCF/Lef:H2B-GFP* reporter mice, anti-β-catenin immunofluorescence was increased in the nuclei of astrocytes in the deprived barrel cortex (Figure 4F-G). Importantly, this induction of canonical Wnt signling in wild-type mice appeared to be astrocyte specific as indicated by increased nuclear β-catenin in nuclei co-labeled with the astrocyte nuclear marker SOX9, but no increase in nuclear β-catenin in non-astrocyte, SOX9-negative nuclei in the deprived barrel cortex after whisker lesioning (Figure 4F, H).

**Figure 4.**
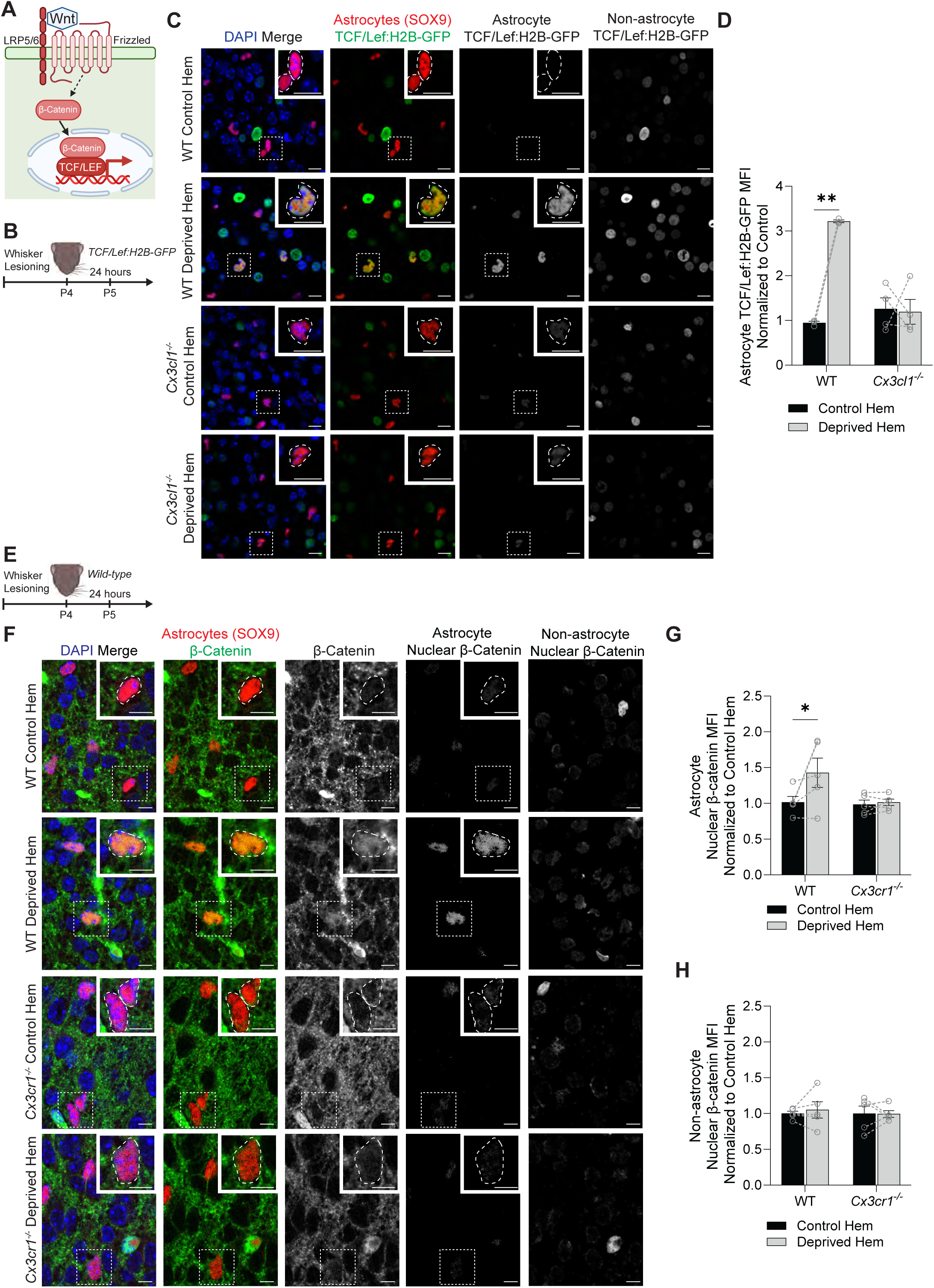
Increased Wnt receptor signaling in layer IV astrocytes is CX3CL1/CX3CR1-dependent. (A) Diagram of the canonical Wnt siganling pathway in which secreted Wnt ligands bind to frizzled and LRP5/6 receptors on the receiving cell, leading to accumulation of β-catenin in the nucleus and activation of TCF/LEF transcription factors. (B) Diagram of experimental protocol used to assess the activity of TCF/LEF transcription factors at 24 hours after whisker lesioning in *TCF/Lef:H2B-GFP* reporter mice. (C) Representative fluorescent images of brain sections containing layer IV of the barrel cortex from deprived and control hemispheres (hem) of wild-type and *Cx3cl1^-/-^* mice at 24 hours after whisker lesioning. Sections were immunolabeled with anti-GFP (green) to assess the activity of the Wnt signaling reporter *TCF/Lef:H2B-GFP*, DAPI (blue) to label nuclei, and anti-SOX9 (red) to label astrocyte nuclei. The right-most panels show anti-GFP immunofluorescence, representing *TCF/Lef:H2B-GFP*, colocalized with astrocytes (overlapping with anti-SOX9) and with non-astrocytes (overlapping with DAPI but not anti-SOX9). Insets depict SOX9^+^ astrocyte nuclei, indicated by white dotted outlines. Scale bars 10 µm. (D) Quantification of the average *TCF/Lef:H2B-GFP* mean fluorescence intensity (MFI) in SOX9^+^ astrocytes shows increased Wnt signaling reporter activity in the deprived hem of wild-type mice, but no difference in *Cx3cl1^-/-^* mice (Repeated measures 2-way ANOVA with Sidak’s post-hoc test: *n* = 3 wild-type mice, 4 *Cx3cl1^-/-^* mice*^-^.* ** *p* < 0.01). (E) Diagram of experimental protocol used to assess levels of nuclear β-catenin at 24 hours after whisker lesioning. (F) Representative fluorescent images of brain sections containing layer IV of the barrel cortex from deprived and control hem of wild-type and *Cx3cr1^-/-^* mice at 24 hours after whisker lesioning. Sections were immunolabeled with anti-SOX9 (red) to label astrocyte nuclei, DAPI (blue) to label nuclei, and anti-β-catenin (green). The two right-most columns show anti-β-catenin in astrocyte nuclei (overlapping with anti-SOX9), and anti-β-catenin in the nuclei of non-astrocytes (overlapping with DAPI but not anti-SOX9). Insets depict SOX9^+^ astrocyte nuclei, indicated by white dotted outlines. Scale bars 10 µm. (G) Quantification of the average anti-β-catenin MFI in SOX9^+^ astrocyte nuclei shows increased nuclear β-catenin in the deprived hem of wild-type mice, but no difference in *Cx3cr1^-/-^* mice (Repeated measures 2-way ANOVA with Sidak’s post-hoc test: *n* = 5 wild-type mice, 5 *Cx3cr1^-/-^* mice. * *p* < 0.05). (H) Quantification of the average anti-β-catenin MFI in the nuclei of non-astrocytes (SOX9-negative) shows no difference between the control and deprived him in both wild-type and *Cx3cr1^-/-^* mice (Repeated measures 2-way ANOVA with Sidak’s post-hoc test: *n* = 5 wild-type mice, 5 *Cx3cr1^-/-^* mice). All data are presented as mean ± SEM.

Using these same assays, we also tested whether Wnt receptor activation in astrocytes is dependent on CX3CL1/CX3CR1 signaling. Because *Cx3cr1^-/-^* mice also express GFP, we analyzed *TCF/Lef:H2B-GFP* reporter activity in *Cx3cl1^-/-^* mice crossed with *TCF/Lef:H2B-GFP* mice and found that the increase in reporter activity in astrocytes following whisker lesioning was abolished in these knockout mice (Figure 4C-D). We further assessed the role of the receptor CX3CR1 by measuring levels of nuclear astrocytic β-catenin in *Cx3cr1^-/-^* (*Cx3cr1^EGFP/EGFP^*) mice. Similar to *Cx3cl1^-/-^* mice, induction of nuclear β-catenin in astrocytes was blocked in *Cx3cr1^-/-^* mice following whisker lesioning (Figure 4F-G). Together, these data demonstrate that whisker lesioning in P4 mice induces upregulation of canonical Wnt signaling specifically in layer IV astrocytes within the barrel cortex. These data further demonstrate that Wnt signaling in astrocytes is induced in a microglial CX3CL1-CX3CR1 signaling-dependent manner.

### Microglial Wnt release is required to induce decreased astrocyte-synapse interactions and synapse remodeling

Our data demonstrate that CX3CL1-CX3CR1 signaling is critical to induce Wnt receptor signaling in astrocytes and to induce astrocytes to reduce their contact with synapses in response to whisker lesioning in P4 mice. However, it was unclear if microglial Wnt release was necessary for astrocytes to alter their synaptic contacts or if a different CX3CL1-CX3CR1-dependent signal caused the astrocyte process retraction in parallel. To directly interrogate these possibilities, we leveraged pharmacological and cell type-specific molecular approaches. To genetically block Wnt release from microglia we leveraged mice harboring floxed alleles of wntless (*Wls*),^33^ which is required for trafficking and secretion of Wnt proteins from cells (Figure 5A).^34–36^ Given the narrow time window for our experiments, gene deletion approaches using the inducible form of CreER in microglia^37^ were not compatible with our experiments. Therefore, we chose to specifically ablate *Wls* in microglia using a BAC transgenic *Cx3cr1^Cre^* line,^38, 39^ in which *Cx3cr1^Cre^* is randomly inserted into the genome such that endogenous *Cx3cr1* expression is not impacted. We refer to these mice as *Wls^F/F^*; *Cx3cr1^Cre^* (WLS cKO) mice (Figure 5B). We note that, in addition to microglia, CX3CR1 is also expressed by other brain border macrophages and peripheral immune cells at this age. However, our analyses are all performed in the brain parenchyma within the barrel cortex. In this region and context, microglia are the only CX3CR1 expressing cells. Deletion of *Wls* specifically in microglia at this age was confirmed by *in situ* hybridization (Figure S3). We then assessed synapse remodeling following whisker lesioning at P4 and we found that preventing Wnt release from microglia was sufficient to block loss of VGluT2^+^ TC presynaptic terminals in layer IV in the deprived hemisphere at 6 days post-lesioning (Figure 5C-D). Importantly, this phenotype was also found in postnatal mice treated daily from P3-P9 with XAV939,^40^ a brain-permeable pharmacological inhibitor of canonical Wnt signaling (Figure S4).

**Figure 5.**
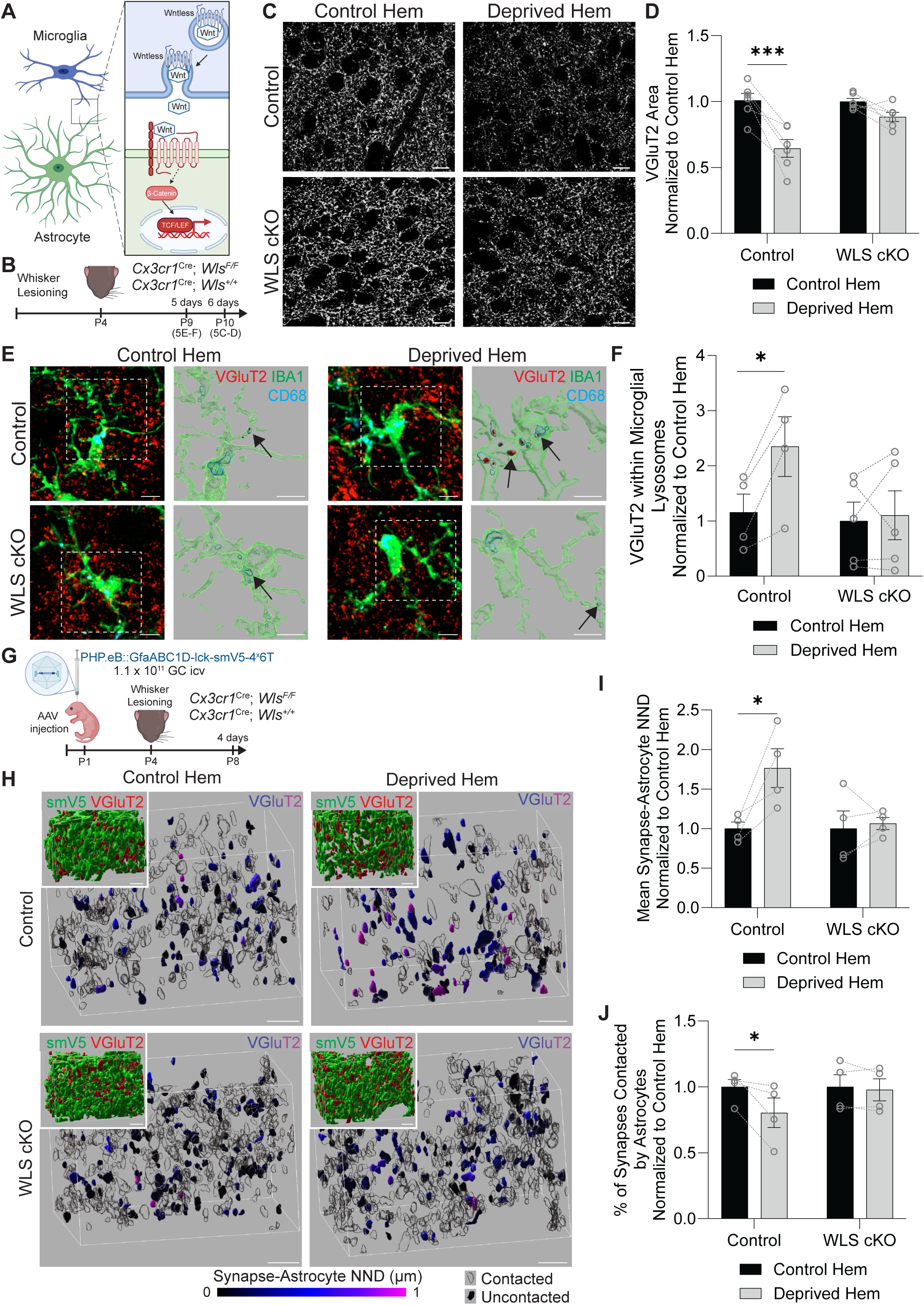
Microglial Wnt release is required to induce decreased astrocyte-synapse interactions and synapse remodeling. (A) Diagram of microglia-astrocyte Wnt signaling. Wnt ligand is bound to the trafficking protein wntless (WLS) in microglia and then secreted. Secreted Wnt ligand can bind frizzled and LRP5/6 receptors to stimulate canonical Wnt receptor signaling in astrocytes, which involves accumulation of β-catenin in the nucleus and activation of TCF/LEF transcription factors. (B) Diagram of experimental protocol used to assess the density of VGluT2^+^ synaptic terminals in layer IV of the barrel cortex at 6 days after whisker lesioning or microglial engulfment of VGluT2^+^ synaptic terminals at 5 days after whisker lesioning in *Cx3cr1^Cre^*; *Wls^F/F^* (WLS cKO) and *Cx3cr1^Cre^*; *Wls^+/+^*(Control) mice. (C) Representative fluorescent images of anti-VGluT2^+^ thalamocortical synaptic terminals in layer IV of the barrel cortex in the control and deprived hemispheres (hem) at 6 days post-lesioning in WLS cKO and control mice. Scale bars 10 µm. (D) Quantification of the density of anti-VGluT2^+^ thalamocortical synaptic terminals in layer IV of the barrel cortex at 6 days post-lesioning shows reduced density in the deprived hem compared to the control hem in control mice, but not WLS cKO mice (Repeated measures 2-way ANOVA with Sidak’s post hoc test: *n* = 6 control mice, 6 WLS cKO mice. *** *p* < 0.001). (E) Representative (left) fluorescent max intensity projection images and (right) Imaris surface reconstructions of anti-VGluT2^+^ thalamocortical synaptic terminals (red), anti-CD68^+^ lysosomes (cyan), and anti-IBA1^+^ microglia (green) in layer IV of the barrel cortex in the control and deprived hems of WLS cKO and control mice at 5 days post-lesioning. Imaris reconstructions are from white outlined regions of fluorescent images and depict only the anti-VGluT2^+^ synaptic material contained within anti-CD68^+^ lysosomes inside anti-IBA1^+^ microglia (examples indicated by black arrows). Scale bars 5 µm. (F) Quantification of anti-VGluT2^+^ synaptic material within microglial lysosomes at 5 days after whisker lesioning shows more engulfed anti-VGluT2^+^ material in the deprived hem of the barrel cortex compared to the control hem in control mice, but not in WLS cKO mice (Repeated measures 2-way ANOVA with Sidak’s post-hoc test: *n* = 4 control mice, 5 WLS cKO mice. * *p* < 0.05). (G) Diagram of experimental protocol used to assess astrocyte-synapse interactions in WLS cKO and control mice at 4 days after unilateral whisker lesioning. Astrocytes were labeled with membrane-bound V5 protein by intracerebroventricular (icv) injection of PHP.EB::GfaABC1D-lck-smV5-4^x^6T adeno-associated virus (AAV) at P1. (H) Representative Imaris surface reconstructions of anti-VGluT2^+^ thalamocortical synaptic terminals in layer IV barrel cortex from the control and deprived hems of WLS cKO and control mice at 4 days after unilateral whisker lesioning. Anti-VGluT2^+^ thalamocortical synaptic terminals are pseudo-colored by the nearest neighbor distance (NND; corrected for expansion index) to the nearest astrocyte process. Anti-VGluT2^+^ surfaces with NND of 0 are considered “contacted” and are shown in transparent black. Insets: anti-GFP^+^ astrocytes (green) and anti-VGluT2^+^ thalamocortical synaptic terminals (red). Scale bars 2 µm (corrected for expansion index). (I) Quantification of the mean synapse-astrocyte NND in layer IV of the barrel cortex at 4 days after unilateral whisker lesioning shows increased NND in the deprived hem of control mice compared to the control hem, but not in WLS cKO mice (Repeated measures 2-way ANOVA with Sidak’s post-hoc test: *n* = 4 control mice, 4 WLS cKO mice. * *p* < 0.05). (J) Quantification of the percentage of synapses contacted by astrocytes in layer IV of the barrel cortex at 4 days after unilateral whisker lesioning shows reduced astrocyte-synapse contact in the deprived hem of control mice compared to the control hem, but not in WLS cKO mice (Repeated measures 2-way ANOVA with Sidak’s post-hoc test: *n* = 4 control mice, 4 WLS cKO mice. * *p* < 0.05). All data are presented as mean ± SEM. See also Figures S3, S4, and S5.

We next assessed microglia and astrocyte interactions with synapses. We first found that microglial engulfment of VGluT2^+^ synaptic material was blocked in WLS cKO mice at 5 days post-lesioning (Figure 5E-F). We then assessed astrocytes. For these experiments, WLS cKO mice were intracerebroventricularly injected at P1 with PHP.eB::GfaABC1D-lck-smV5-4x6T adeno-associated virus (AAV)^41^ to label astrocytes with membrane bound smV5 (Figure S5A-C). In WLS cKO mice, there was no longer a change in astrocyte process density (Figure S5D-F) or astrocyte association with synapses (Figure 5G-J) in layer IV of the barrel cortex at 4 days after whisker lesioning. Together, these data show that microglial Wnt release is sufficient to induce astrocyte process retraction, microglial synapse engulfment, and TC input elimination in response to whisker lesioning. Furthermore, in conjunction with our previous data showing that astrocyte Wnt receptor signaling and astrocyte process retraction are dependent on CX3CL1-CX3CR1 signaling, they suggest that activation of CX3CR1 in microglia leads to synapse elimination by regulating Wnt production and/or release from microglia to astrocytes.

## Discussion

It is becoming increasingly appreciated that both microglia and astrocytes can carry out synapse engulfment and pruning in the developing brain.^7^ In some cases, this is within the same brain region.^2^ This begs the question of whether these two cell types are communicating to carry out this critical function for sculpting and remodeling neural circuits. Here, we show that microglia communicate directly with astrocytes through Wnt release to stimulate astrocytes to remodel their processes and subsequently permit microglia-mediated synapse removal. In addition, numerous studies have shown a role for CX3CL1-CX3CR1 signaling in microglia-mediated regulation of synapses,^14–17^ but mechanisms downstream of this ligand-receptor signaling have remained elusive. Our study provides insight into how CX3CL1-CX3CR1 signaling is regulating microglia-mediated synapse remodeling. In this context, CX3CL1-CX3CR1 activation does not induce classical chemokine signaling but instead engages activity-dependent microglia-to-astrocyte communication via Wnts prior to microglia-mediated synapse engulfment. Wnt release from microglia engages astrocytes to then redistribute their processes further away from synapses, which is subsequently permissive for microglia to engulf and remove synapses.

Previously, it has been shown that both microglia and astrocytes can similarly engulf and remove synapses. However, it has been unclear to what extent there is coordination between these two cell types. In the mouse cortex, it has been shown that microglia and astrocytes engulf separate domains of the cell during apoptotic cell death with microglia engulfing the cell body while the astrocytes engulf the processes.^42^ Also, in the mouse spinal cord and thalamus, astrocyte secretion of IL-33 stimulates microglia to upregulate phagocytic pathways to engulf and remove synapses.^4–6^ These studies indicate intercellular crosstalk between astrocytes and microglia occurs to coordinate synapse engulfment. However, the “eat-me” signals thought to tag synapses for elimination (e.g. C1q and Gas6) can be bound by phagocytic receptors expressed by both cell types, astrocytes via MERTK and MEGF10 and microglia via C3R, suggesting the potential for competition. Also, in disease contexts, several examples of microglia-astrocyte crosstalk have been identified.^43^ For example, in a mouse model of neuroinflammatory neurodegeneration, TGFα and VEGF-B produced by microglia bind respectively to ErbB1 and FLT1 in astrocytes and modulate astrocytes.^44^ Nevertheless, during physiological settings, it remains unclear if microglia can communicate with astrocytes to regulate synapse remodeling and whether this crosstalk is activity-dependent. Here, we show that CX3CR1-dependent Wnt release from microglia is an activity-dependent mechanism by which microglia signal to astrocytes to facilitate synapse removal by microglia.

In the current study, we identified that Wnt release from microglia induced Wnt receptor signaling in astrocytes to induce astrocyte process remodeling away from synapses. Wnt signaling is a known regulator of cell morphology and the actin cytoskeleton, including in astrocytes.^32^ Although not directly tested, we suspect that the reduction in astrocyte-synapse interactions is caused by Wnt receptor signaling induced changes in the astrocyte actin cytoskeleton, which then leads to decreased astrocyte process density and reduced contact with synapses. The precise Wnt(s) released by microglia and how CX3CR1 signaling regulates this release also remain open questions. One possibility is through Ca^2+-^dependent Wnt release from microglia downstream of CX3CR1 activation. CX3CR1 is a GPCR coupled to Gi and Gz proteins, which can induce release of Ca^2+^ from internal stores.^45^ It is possible that Wnt release from microglia is Ca^2+^ dependent. Also, it is noted that in addition to microglia, CX3CR1 is also expressed by brain border macrophages and peripheral immune cells. In disease contexts, peripheral-derived macrophages can communicate with astrocytes.^46^ However, in the context of layer IV of the developing barrel cortex where the blood brain barrier is intact, microglia are the only source of CX3CR1. Thus, our results are most consistent with a microglia-dependent mechanism.

Our observation that astrocytes decrease contact with synapses prior to their removal by microglia suggests that reduced astrocyte-synapse contact is permissive for microglia-mediated synapse removal. However, there are multiple potential mechanisms by which reduced astrocyte-synapse contact could facilitate microglia-mediated synapse removal. One possibility is that astrocytes might physically shield synapses from removal. Microglia are known to remove specific subsets of synapses by recognizing “eat-me” signals such as C1q and/or phosphatidyl serine that “tag” synapses for removal,^7^ requiring direct contact between microglia and synapses for their removal. Reductions in astrocyte-synapse contact could allow for increased contact between microglia and synapses, allowing greater access to these “eat me” signals. Astrocyte perisynaptic processes also perform important metabolic, trophic, and signaling functions,^47^ which are likely altered as astrocyte-synapse contact is reduced. Synapses without astrocyte contact might have long term depression, reduced synapse stability, or impaired synapse maturation, leading to their weakening and eventual removal by microglia. Indeed, one study genetically reduced astrocyte-synapse contacts via CRISPR/Cas9 depletion of ezrin in astrocytes.^48^ This manipulation led to increased glutamate spillover and increased NMDA receptor activation in the hippocampus, reinforcing the link between astrocyte-synapse contact and synaptic function. Importantly, these two alternatives are not mutually exclusive. Reduced astrocyte-synapse contact could weaken synapses, leading to increased surface-tagging of synapses with “eat me” signals while also allowing microglia greater access to contact and remove synapses.

Due its accessibility and the wealth of knowledge on its circuit function and development, the barrel cortex is an ideal circuit to work out cellular/molecular mechanisms of synapse remodeling.^7^ Using this circuit, we uncovered the involvement of microglia-astrocyte Wnt signaling and the retraction of astrocyte processes from synapses in the regulation of microglia-mediated synapse remodeling in response to changes in neuronal activity. One intriguing possibility is that this could be a broader mechanism by which microglia and astrocytes coordinate to remove synapses in health and disease. Indeed, by comparing our results to other transcriptomic studies in astrocytes, we found evidence that Wnt receptor signaling is activated in astrocytes in other contexts such as Alzheimer’s disease and epilepsy in which synapse loss and astrocyte morphological changes are known to occur. These data match a recent comparison of reactive astrocytes showing that β-catenin, the main effector of the Wnt signaling pathway, is among the core transcriptional regulators of astrocyte reactivity shared across divergent conditions.^49^ Also, among the defining features of reactive astrocytes are morphological changes,^50^ which may affect their association with synapses. Further evidence of the broad applicability of our findings is that WLS, the gene required for Wnt secretion, is among the genes most significantly reduced in microglia in human patients with Autism spectrum disorder^51^ and that the Wnt signaling pathway is a target for neuropsychiatric drugs including lithium, Ritalin, fluoxetine, and clozapine.^52, 53^ Evidence from mouse models also indicates that Wnt ligand expression in microglia can be affected by early life inflammatory insults, another key risk factor for neuropsychiatric disorders.^54^ Thus, microglia-astrocyte Wnt signaling may play an overlooked role in these disorders. Finally, under physiological conditions such as during sleep or learning and memory tasks, electron microscopy studies have shown that astrocytes change their association with synapses concomitant with changes in synapse numbers^25, 48, 55, 56^ and during *in vivo* time lapse imaging experiments, astrocyte-synapse coverage is predictive of synapse stability.^57^ It is, thus, possible that Wnt-dependent microglia-astrocyte interactions also regulate synapse remodeling in these contexts.

In summary, our findings provide deep mechanistic insight into how CX3CR1-activation leads to synapse removal by microglia via Wnt signaling. We further provide data that supports astrocyte process rearrangement away from synapses as a key step in permitting microglia to engulf and remove synapses following sensory deprivation. Reductions in astrocyte-synapse interactions could facilitate microglial synapse removal by increasing synaptic accessibility to microglia and/or weakening synaptic strength to facilitate engulfment of weak synapses by microglia. Our data further show that Wnt receptor signaling in astrocytes occurs in a broad range of physiological disease conditions where changes in astrocyte morphology and synapse elimination have been documented. Further studies will be needed to test how and in which contexts changes in astrocyte-synapse contacts are induced by microglia-mediated release of Wnts, leading to subsequent synapse removal by microglia.

## Acknowledgements

This work was supported by NIMH-R01MH113743 (DPS), NIMH-R01MH118329 (AS), NINDS-R01NS117533 (DPS), NINDS-R01NS106721 (AS), NIA-RF1AG068281(DPS), NIA-RF1AG068558 (AS), NIA-R01AG072489 (AS), ERC-951515 (AS), Robin Chemers Neustein Postdoctoral Award (PA), Alzheimer’s Association AARG-22-974642 (PA), Alfred P. Sloan Foundation JFRASE (PA), NIH T32AG049688 (AB), Leon Levy Scholarship in Neuroscience (AB), NINDS-F31NS117053 (GG), and the Dr. Miriam and Sheldon G. Adelson Medical Research Foundation (DPS). Schematics were created with Biorender.com. We thank Rockefeller Genomics Core for quality control and sequencing libraries and Y-H. E. Loh for TRAP sequencing bioinformatics analysis. We thank Dr. Sergio Lira for providing *Cx3cl1^-/-^* mice, Dr. David Rowitch for providing *Wls^Flox^* mice, Dr. Ben Barres for providing *Aldh1l1^EGFP^* mice, and Dr. Staci Bilbo for providing *Cx3cr1^Cre^* mice. We thank Dr. Lukasz Szewczyk and Dr. Marta Wisniewska for providing protocols for immunofluorescent detection of nuclear β-catenin. We thank Gregory Hendricks and the UMass Chan Electron Microscopy Facility for help staining and imaging samples for electron microscopy. We thank Ugur Celik and the UMass Chan 3D printing core for producing custom molds for cortical flatmount preparations. We also thank Shannon Becker for technical assistance.

## Author Contributions

TEF and DPS designed the study. TEF and DPS wrote the manuscript. TEF performed most experiments and analyzed most data. YL performed viral injections, assisted in expansion microscopy imaging and analysis, and performed experiments to analyze astrocyte cell body density. CO performed experiments to assess synapse density in WLS conditional knockout mice and mice treated with XAV939. MAB assisted in analysis of synapse density in wild-type mice and analysis of synapse engulfment in wild-type and WLS conditional knockout mice. GG, AB, PA, and AS performed the TRAP-Seq experiments and assisted in initial data analysis. AS provided critical input into study design and feedback on writing of the manuscript.

## Inclusion and Diversity

We worked to ensure sex balance in the selection of non-human subjects. We worked to ensure diversity in experimental samples through the selection of the genomic datasets. One or more of the authors of this paper self-identifies as a gender minority in their field of research. While citing references scientifically relevant for this work, we also actively worked to promote gender balance in our reference list. We avoided “helicopter science” practices by including the participating local contributors from the region where we conducted the research as authors on the paper.

## STAR Methods

### Resource Availability

#### Lead contact

Further information and requests for resources should be directed to and will be fulfilled by the lead contact, Dorothy Schafer.

#### Materials availability

This study did not generate new unique reagents.

#### Data and code availability

TRAP-seq data have been deposited at GEO and are publicly available as of the date of publication. Any additional information required to reanalyze the data reported in this paper is available from the lead contact upon request.

### Experimental model and subject details

Wild-type C57Bl6/J (Stock #000664), *Rosa26^mTmG^* (Stock# 007676, B6.129(Cg)-*Gt(ROSA)^26Sortm4(ACTB-tdTomato,-EGFP)Luo^*/J),^18^ *Cx3cr1^EGFP^* (Stock# 005582, B6.129P2(Cg)-*Cx3cr1^tm1Litt^*/J),^20^ and *TCF/Lef:H2B-GFP* (Stock# 013752, STOCK Tg(TCF/Lef1-HIST1H2BB/EGFP)61Hadj/J)^30^ mice were obtained from Jackson Laboratories (Bar Harbor, ME). *Aldh1l1^CreER^*mice (Stock# 031008, B6N.FVB-Tg(Aldh1l1-cre/ERT2)1Khakh/J)^19^ were obtained from Jackson Laboratories (Bar Harbor, ME) and backcrossed over 12 generations on a C57Bl6/J (Jackson Laboratories) background. *Cx3cl1^-/-^* (*Cx3cl1^tm1Lira^*) mice^21^ were provided by Dr. Sergio Lira (Ichan School of Medicine, Mount Sinai). *Wls^Flox^* (129S-*Wls^tm1.1Lan/J^*) mice^33^ were provided by Dr. David Rowitch (University of California, San Francisco). *Aldh1l1^EGFP^* (STOCK Tg(Aldh1l1-EGFP)OFC789Gsat/Mmucd) mice^58^ on a C57Bl6 background were provided by Dr. Ben Barres (Stanford University).^59^ *Cx3cr1*^Cre^ (STOCK Tg(Cx3cr1-cre)MW126Gsat/Mmcd) mice^38^ on a C57Bl6/N background were provided by Dr. Staci Bilbo (Harvard University)^39^ and subsequently backcrossed over 12 generations on a C57Bl6/J (Jackson Laboratories) background. *Cx3cr1^CreER/+ (Litt)^* (B6.129P2(Cg)-*Cx3cr1^tm2.1(cre/ERT2)Litt^*/WganJ) mice^60^ were a generous gift from Dr. Dan Littman (New York University). *Eef1a1^LSL.eGFPL10a/+^* (B6.Cg-*Eef1a1^tm1Rck^*/J) mice^61^ were generously provided by Dr. Ana Domingos (Instituto Gulbenkian de Ciência, PT) and Dr. Jeff Friedman (Rockefeller University). Aldh1l1^EGFP-L10a^ (B6;FVB-Tg(Aldh1l1-EGFP/Rpl10a)JD130Htz/J) mice^22^ were generously provided by Dr. Nathaniel Heintz (Rockefeller University). All mice were maintained on a C57Bl6/J background. Animals were maintained on a 12 hour light/dark cycle with food and water provided ad libitum. All animals were healthy and were not immune compromised. Experimental mice were not involved in previous procedures or studies. Littermates of the same sex were randomly assigned to experimental groups. Both males and females were used for all experiments. Experiments were performed in mice aged 1-10 days old. The specific ages used for each experiment are indicated in the figures. All animal experiments were performed in accordance with Animal Care and Use Committees (IACUC) and under NIH guidelines for proper animal use and welfare.

### Method Details

#### Whisker lesioning

Whisker lesioning was performed as described previously.^13^ Briefly, whiskers of anesthetized P4 mice were unilaterally lesioned by cauterization of the right whisker pad with a high temperature cautery (Bovie Medical Corporation; Clearwater, FL). Care was taken to ensure that all whiskers on that side were lesioned.

#### Immunohistochemistry

Mice were anesthetized and transcardially perfused with 0.1 M phosphate buffer (PB), followed by 4% paraformaldehyde (PFA, Electron Microscopy Sciences; Hatfield, Pennsylvania) in 0.1 M PB. For analyses in layer IV of the barrel cortex, cortical flatmounts were prepared as described previously in order to visualize the entire barrel field in a single section.^13^ Briefly, the cortex from the right and left hemisphere was dissected away from the midbrain, placed in open-face custom molds with a 250 µm-thick chamber 3D-printed using ABS-M30 material (Stratasys Inc.; Eden Praire, Minnesota), submerged in 4% PFA solution, covered with a glass slide to gently flatten, and post-fixed overnight at 4°C. For analyses in cortical sections, whole brain tissue was post-fixed overnight at 4°C in 4% PFA solution. The following day, tissue was washed three times in 0.1 M PB before storing in 30% sucrose in 0.1 M PB. 40 µm thick sections were cut using a microtome or a vibratome. They were subsequently blocked for 1 hour in 0.1 M PB containing 10% normal goat serum (NGS; G9023 Sigma-Aldrich; St. Louis, MO) and 0.3% Triton X-100 (X100 Sigma-Aldrich) (PBTGS). Blocked sections were then incubated with primary antibodies in PBTGS overnight at room temperature, washed 3 times with 0.1 M PB, incubated for 2-3 hours with secondary antibodies in PBTGS at room temperature, washed 3 times with 0.1 M PB, and mounted on glass slides using Fluoroshield mounting media containing DAPI (F6057 Sigma-Aldrich). For immunofluorescent analysis of TCF/Lef:H2B-GFP tissue, sections were stained with Hoechst 33342 dye (H3570 Thermo Fisher Scientific; Waltham, MA) and mounted using CitiFluor mounting medium (CFM-3 Electron Microscopy Sciences).

For immunofluorescent analysis of β-catenin, sucrose-incubated whole brain tissue was cryo-embedded in a 2:1 mixture of 30% sucrose and Tissue-Tek OCT (Sakura Finetek; Torrance, California), frozen on dry-ice and stored at −80°C. 20 µm coronal sections containing whisker barrels were mounted on slides, washed 3 times in phosphate buffered saline (PBS) containing 0.2% Triton-X (PBST), and incubated in Liberate Antibody Binding Solution (24310 Polysciences, Inc; Warrington, PA) overnight to quench endogenous *Cx3cr1^EGFP^* signal. Sections were then washed 3 times with PBST, placed in 10 mM sodium citrate pH 6.0 with 0.05% Tween warmed to 95°C for 40 min for antigen retrieval. Following antigen retrieval, sections were washed once with PBST, blocked for 1 hour in 5% normal donkey serum (NDS) with 0.3 M glycine in PBS and incubated overnight at 4°C with primary antibodies in 1% NDS in PBS. After 3 washes in PBST, sections were then incubated with secondary antibodies raised in donkey in 1% NDS in PBS for 1 hour at room temperature, washed 3 times with PBST, and mounted with Fluoroshield mounting media containing DAPI (F6057 Sigma-Aldrich).

Primary antibodies used: guinea pig anti-VGluT2 (1:2000 for synapse area; 1:1000 for synapse engulfment; 135 404 Synaptic Systems; Göttingen, Germany), chicken anti-GFP (1:1000; ab13970 Abcam; Cambridge, United Kingdom), rat anti-CD68 (1:200; MCA1957 Bio-Rad; Hercules, CA), rat anti-LAMP2 (1:200; ab13524 Abcam), rabbit anti-SOX9 (1:500; ab185966 Abcam), mouse anti-β-catenin Alexa Fluor 647 (1:100; sc-7963 AF647 Santa Cruz Biotechnology; Dallas, TX), chicken anti-IBA1 (1:500; 234 009 Synapatic Systems), rat anti-V5 (1:100; Ab00136-6.1 Absolute Antibody; Boston, MA). Secondary antibodies were used at 1:1000 dilution: goat anti-chicken IgY Alexa Fluor 488 (A-11039 Thermo Fisher Scientific), goat anti-guinea pig IgG Alexa Fluor 594 (A-11076 Thermo Fisher Scientific), goat anti-rat IgG Alexa Fluor 647 (A-21247 Thermo Fisher Scientific), goat anti-rabbit IgG Alexa Fluor 647 (A-21245 Thermo Fisher Scientific), goat anti-rabbit IgG Alexa Fluor 488 (A-11034 Thermo Fisher Scientific), goat anti-rat IgG Alexa Fluor 488 (A-11006 Thermo Fisher Scientific), donkey anti-guinea pig IgG Alexa Fluor 488 (706-545-148 Jackson ImmunoResearch Labs; West Grove, PA), donkey anti-rabbit IgG Alexa Fluor 594 (A-21207 Thermo Fisher Scientific).

#### Synapse density analysis

Analysis of synapse density was performed as previously described.^13^ Briefly, fluorescent immunolabeled 40 µm thick brain sections were imaged at 63x on a Zeiss LSM700 scanning confocal microscope equipped with 405nm, 488nm, 555nm, and 639nm lasers and ZEN acquisition software (Zeiss; Oberkochen, Germany). For each animal, single plane images centered within a barrel in layer IV of the barrel cortex were acquired from 4 fields of view (FOV) per hemisphere. Images were analyzed in FIJI software by experimenters blind to the identity of the samples.^62^ After background subtraction (rolling ball radius 10 pixels) and despeckling, the VGluT2^+^ signal was manually thresholded and the Analyze Particles function (particle size 0.2 pixels to infinity; ImageJ plugin, NIH) was used to measure the total area of VGluT2^+^ presynaptic terminals. Data for each hemisphere per animal were averaged across the 4 FOV.

#### Engulfment analysis

Analysis of engulfment of VGluT2^+^ material was performed as previously described.^3,13, 63^ Briefly, fluorescent immunolabeled 40 µm thick brain sections were imaged at 63x on a Zeiss Observer Spinning Disk Confocal microscope equipped with diode lasers (405nm, 488nm, 594nm, 647nm) and ZEN acquisition software (Zeiss). For each animal, z-stack images (0.22 µm step size) centered within a barrel in layer IV of the barrel cortex were acquired from 6 to 8 FOV per hemisphere for microglia and 4 FOV for astrocytes. Images were background subtracted (rolling ball radius 50 pixels for microglia and astrocytes, 10 pixels for CD68 and LAMP2, 10 pixels for VGluT2) using FIJI software, then processed in Imaris (Bitplane; Zurich, Switzerland) as previously described.^3, 13, 63^ Data analysis was performed by experimenters blind to the identity of the samples. For engulfment analysis in microglia, data for each hemisphere per animal was averaged across 15-20 individual cells per animal. For engulfment analysis in astrocytes, since it was difficult to determine the boundaries of individual astrocytes, surface renderings were generated for all astrocytes within the barrel region of each z-stack image. Engulfment analysis was restricted to anti-VGluT2^+^ material contained within CD68^+^ lysosomes for microglia and LAMP2^+^ lysosomes for astrocytes. Data for each hemisphere per animal were averaged across the 4 FOV.

#### Tamoxifen injections

For expansion microscopy experiments (see Expansion microscopy), *Aldh1l1^CreER^*; *Rosa26^mTmG/+^* mice were injected once daily intraperitoneally (i.p.) with 50 µg tamoxifen (T5648 Sigma-Aldrich) dissolved in corn oil at 1 mg/mL at ages P1, P2, P3, and P4. For TRAP-Seq experiments (see TRAP-Seq), *Cx3cr1^CreER/+ (Litt)^*; *Eef1a1^LSL.eGFPL10a/+^*mice were orally administered 50 µg tamoxifen (10 mg/mL in corn oil) once daily at ages P1, P2, and P3.

#### Expansion microscopy

For expansion microscopy, 40 µm tangential brain sections were prepared as described above (see Immunohistochemistry). Sections were incubated overnight in Liberate Antibody Binding Solution (24310 Polysciences, Inc) for antigen retrieval and to eliminate endogenous mTomato fluorescence in *Rosa26^mTmG^* mice, then washed three times in 0.1 M PB. Sections were then blocked for 4 hours in PBTGS or 0.1 M PB with 10% NDS and 0.3% Triton-X (PBTDS) and incubated 3 days overnight at 4°C with primary antibody diluted in PBTGS or PBTDS. Following primary antibody incubation, sections were washed three times in 0.1 M PB, incubated overnight at room temperature with secondary antibody diluted in PBTGS or PBTDS, and washed 3 times in 0.1 M PB.

Primary antibodies used: guinea pig anti-VGluT2 (1:1500; 135 404 Synaptic Systems), chicken anti-GFP (1:1000; ab13970 Abcam), rat anti-V5 (1:100; Ab00136-6.1 Absolute Antibody), rabbit anti-ALDH1L1 (1:500; 85828 Cell Signaling Technology; Danvers, MA). Secondary antibodies were used at 1:300 dilution: goat anti-chicken IgY Alexa Fluor 488 (A-11039 Thermo Fisher Scientific), goat anti-guinea pig IgG Alexa Fluor 594 (A-11076 Thermo Fisher Scientific), goat anti-rat IgG Alexa Fluor 594 (A-11007 Thermo Fisher Scientific), goat anti-guinea pig IgG Alexa Fluor 488 (A-11073 Thermo Fisher Scientific), goat anti-rabbit IgG, Atto 647N (40839 Sigma-Aldrich), donkey anti-guinea pig IgG Alexa Fluor 488 (706-545-148 Jackson ImmunoResearch), donkey anti-rat IgG Alexa Fluor 594 (A-21209 Thermo Fisher Scientific).

Prior to gel embedding and expansion, pre-expansion images of the barrel cortex region of immunolabeled brain sections were acquired at 10x and 20x on a Zeiss Observer Spinning Disk Confocal microscope equipped with diode lasers (405nm, 488nm, 594nm, 647nm) and ZEN acquisition software (Zeiss). Sections were then embedded in polyacrylamide gels and expanded in de-ionized H_2_O as previously described.^64, 65^ Briefly, brain sections were incubated overnight at room temperature in PBS containing 0.01% acryloyl-X SE (A20770 Thermo Fisher Scientific), washed twice with PBS for 15 minutes, and incubated on ice for 30 minutes in gelling solution [0.2% tetramethylethylenediamine (T9281 Sigma-Aldrich), 0.2% ammonium persulfate (A3678 Sigma-Aldrich), 0.01% 4-hydroxy-TEMPO (176141 Sigma-Aldrich), 8.11% sodium acrylate (QC-1489 Combi Blocks; San Diego, CA), 2.35% acrylamide (A9099 Sigma-Aldrich), 0.14% N,N′-methylenebisacrylamide (M7279 Sigma-Aldrich), 10.987% NaCl (S271 Sigma-Aldrich) in PBS]. Gelation chambers were then constructed by transferring brain sections onto glass slides (one section per slide), flanking the section with two stacks of two #1.5 glass coverslips (∼170 µm thick per coverslip), covering the sections with gelling solution, and topping with a glass slide wrapped in parafilm. Completed gelation chambers were incubated for 2 hours at 37°C to polymerize the gels. Polymerized gels were then removed from the gelation chambers, trimmed down with a razor blade, and digested overnight at room temperature with 8 units/mL Proteinase K (P8107S New England BioLabs; Ipswich, MA) in buffer containing 0.5% Triton X-100 (X100 Sigma-Aldrich), 2 mM ethylenediaminetetraacetic acid (EDTA), 50 mM Tris, and 0.8 M NaCl. Digested gels were either expanded for imaging or washed 3 times for 20 minutes with PBS and stored at 4°C for up to a week.

Gels were expanded for imaging by washing 3 times for 20 minutes with de-ionized H_2_O, including a 20 minute incubation in Hoechst 33342 dye (H3570 Thermo Fisher Scientific) during the middle wash. Gels were further trimmed down to the region of interest using a razor blade and transferred to a 6-well plate with a #1.5 cover glass bottom (P061.5HN Cellvis; Burlington, Ontario) pre-coated with 0.1% poly-L-lysine (P8920 Sigma Aldrich) for imaging on a Zeiss Observer Spinning Disk Confocal microscope equipped with diode lasers (405nm, 488nm, 594nm, 647nm), inverted objective lenses, and ZEN acquisition software (Zeiss). Post-expansion images of each gel were acquired at 10x in a FOV contained within the FOV of the pre-expansion images of the same gel. The expansion index for each gel was calculated using FIJI software by comparing the distances between landmarks identified in both the pre-expansion and post-expansion image for at least 3 sets of measurements per gel. The expansion index was calculated for each gel by averaging across these measurements for each gel. For data acquisition, z-stack images (0.50 µm step size) centered within a barrel in layer IV of the barrel cortex were acquired from 4 FOV at 40x with a water-immersion objective. Care was taken to avoid unlabeled astrocytes.

For data analysis, z-stack images were background subtracted (rolling ball radius 50) using FIJI software, then normalized by layer in Imaris (Bitplane; Zurich, Switzerland). Astrocytes labeled with membrane-bound GFP in *Aldh1l1^CreER/+^; Rosa26^mTmG/+^* tissue were 3D rendered using the Imaris surface tool with auto thresholding, 0.25 µm smoothing, voxel size > 150, and sphericity < 0.73. Astrocytes labeled with anti-V5 with 3D rendered using the Imaris surface tool using LabKit, a FIJI plugin for machine learning pixel classification.^66^ For each animal, the same LabKit classification settings were used for all z-stack images for both control and deprived hemispheres. Astrocyte process density was determined by dividing the volume of the astrocyte surface rendering by the total volume of the z-stack image. VGluT2^+^ synapses were 3D rendered using the Imaris surface tool with manual thresholding with 4 µm local background subtraction, 0.25 µm smoothing, and voxel size > 50. Synapse-astrocyte nearest neighbor distances (NND) were calculated by Imaris using VGluT2^+^ surfaces less than 60 µm. VGluT2 surfaces with a NND of 0 were considered contacted by astrocytes and VGluT2 surfaces with NND less than 0 were excluded from analysis. For each z-stack image, we calculated both the mean synapse-astrocyte NND and the percentage of synapses contacted by the astrocytes. Data for each hemisphere per animal were averaged across the 4 FOV.

#### Electron microscopy

For electron microscopy, *Rosa26^mTmG/+^*mice were anesthetized on ice and transcardially perfused at a rate of 6-8 mL/min with 37°C 0.1 M PB to remove blood followed by 60-80 mL of 37°C fixative solution containing 2.5% glutaraldehyde (Electron Microscopy Sciences) and 2% PFA (Sigma) in cacodylate buffer (0.1 M sodium cacodylate (Electron Microscopy Sciences), 0.04% CaCl_2_ (Sigma), pH 7.4). Perfused mice were held head-down for 15 minutes, then brains were dissected out and post-fixed in the same fixative at 4°C overnight. After washing the samples with ice-cold cacodylate buffer, brains were sectioned to 200 μm thickness using a vibratome. Coronal brain sections containing layer IV of the barrel cortex region were identified by endogenous *Rosa26^mTmG^* fluorescence using a Zeiss Observer microscope equipped with Zen acquisition software (Zeiss). Layer IV of the left and right barrel cortex regions was dissected out and stored in fixative solution on ice before processing and embedding.

Samples were then post-fixed in 1% osmium tetroxide aqueous solution (Electron Microscopy Sciences) with 2.5% potassium ferrocyanide (Millipore Sigma) at RT for 1 hour. Sections were rinsed in water and then maleate buffer (Millipore Sigma) (pH=6.0) and stained in 0.05 M maleate buffer containing 1% uranyl acetate (Electron Microscopy Sciences) at 4°C overnight. Sections were dehydrated in series of washes from 5% to 100% ethanol, then two changes of propylene oxide before being infiltrated overnight at room temperature with 1:1 Embed 812 resin: propylene oxide. Finally, sections were embedded in 100% Embed 812 resin between two sheets of Aclar Embedding film (Ted Pella, Inc.) and polymerized at 60°C for 48 h. Once polymerized, the embedding brain sections were mounted on a blank sectioning blocks with a single drop of embedding resin and allowed to polymerize overnight before being trimmed and serial sectioned (60 nm thick). The sections were placed on 200 mesh grids, then stained with 1% uranyl acetate and aqueous lead citrate before being examined using a Tecnai 12 Spirit TEM at 120 kV accelerating voltage. 11500x images were collected using a Gatan Rio9 CCD camera and Gatan DM4 Digital Micrograph software. For each samples, images from 30-50 FOV were collected from the central portion of the section, the portion corresponding to layer IV of the barrel cortex.

Data analysis was performed by experimenters blind to the identity of the samples. In each image, synapses were identified by the presence of an electron-dense synaptic cleft and astrocytes were identified by their irregular shape, electron-sparse cytoplasm, and glycogen granules. For each synapse, it was determined whether there was direct contact with an astrocyte process and the distance was measured from the synaptic cleft to the nearest astrocyte process. Data was collected from 50-70 synapses per sample and averaged for each hemisphere per animal.

#### Astrocyte cell body density analysis

Analysis of astrocyte cell body density was performed using single plane 20x epifluorescence images of fluorescent immunolabeled 40 µm thick brain sections containing the barrel region acquired with a Zeiss Observer microscope equipped with Zen acquisition software (Zeiss). The total number of SOX9^+^ astrocyte nuclei was counted in 4 VGluT2^+^ barrels per hemisphere using FIJI software and divided by the total area of the 4 barrels to determine the astrocyte cell body density.

#### Translating Ribosome Affinity Purification (TRAP) sequencing

Mice were euthanized with CO_2_, and brain regions of interest were dissected and flash-frozen for TRAP. Ribosome-associated mRNA from microglia was isolated from each region as previously described,^67^ where each sample corresponds to a single mouse. Briefly, the brain tissue was thawed in an ice-cold Wheaton 33 low extractable borosilicate glass homogenizer containing 1 mL cell-lysis buffer (20 mM HEPES-KOH, pH 7.3, 150 mM KCl, 10 mM MgCl2, and 1% NP-40, 0.5 mM DTT, 100 μg/ml cycloheximide, and 10 μl/ml rRNasin (Promega; Madison, WI) and Superasin (Thermo Fisher Scientific). After manually homogenizing the samples with 3-5 gentle strokes using the PTFE homogenizer (grinding chamber clearance 0.1 to 0.15mm). Samples were then homogenized in a motor-driven overhead stirrer at 900 r.p.m. with 12 full strokes. The samples were transferred to chilled Eppendorf tubes, and a post-nuclear supernatant was prepared by centrifugation at 4°C, 10 minutes, 2,000 x *g*. To the supernatant, NP-40 (final concentration = 1%) and DHPC (final concentration = 30 mM) were added. Post-mitochondrial supernatant was prepared by centrifugation at 4°C, 10 minutes, 16,000 x *g.* 200 uL Streptavidin MyOne T1 Dynabeads (Thermo Fisher Scientific), conjugated to 1 μg/μl biotinylated Protein L (Pierce Biotechnology; Waltham, MA) and 50 μg each of anti-eGFP antibodies Htz-GFP-19F7 and Htz-GFP-19C8, bioreactor supernatant (Memorial-Sloan Kettering Monoclonal Antibody Facility) were added to each supernatant. After overnight incubation at 4°C with gentle end-over-end rotation, the unbound fraction was collected using a magnetic stand. The polysome-bound beads were washed with high-salt buffer (20 mM HEPES-KOH, pH 7.3, 350 mM KCl, 10 mM MgCl2, 1% NP-40, 0.5 mM DTT, and 100 μg/ml cycloheximide). RNA clean-up from the washed polysome-bound beads and 5% of the unbound fractions was performed using RNeasy Mini Kit (Qiagen; Venlo, Netherlands) following the manufacturer’s instructions. RNA integrity was assayed using an RNA Pico chip on a Bioanalyzer 2100 (Agilent; Santa Clara, CA), and only samples with RIN > 9 were considered for subsequent analysis. Double-stranded cDNA was generated from 1-5 ng of RNA using the Nugen Ovation V2 kit (NuGEN; San Carlos, CA) following the manufacturer’s instructions. Libraries for sequencing were prepared using Nextera XT kit (Illumina; San Diego, CA) following the manufacturer’s instructions. The quality of the libraries was assessed by 2200 TapeStation (Agilent). Multiplexed libraries were directly loaded on NovaSeq (Ilumina) with single-read sequencing for 75 cycles. Raw sequencing data were processed by using Illumina bcl2fastq2 Conversion Software v2.17.

#### Bioinformatic analysis of RNA sequencing datasets

Raw sequencing reads were first quality checked and trimmed using Trim Galore (https://www.bioinformatics.babraham.ac.uk/projects/trim_galore/ v0.6.4; a wrapper program implementing Cutadapt v2.9 https://journal.embnet.org/index.php/embnetjournal/article/view/200 and FastQC v0.11.9 https://www.bioinformatics.babraham.ac.uk/projects/fastqc/) and mapped to the mouse genome (mm10) using the HISAT2 package (v2.2.0).^68^ Reads were counted using featureCounts (v2.0.0) against the Ensembl v99 annotation.^69^ The raw counts were processed through a variance stabilizing transformation (VST) procedure using the DESeq2 package^70^ to obtain transformed values more suitable than the raw read counts for certain data mining tasks. All pairwise comparisons were performed on the count data of entire gene transcripts using the DESeq2 package (v1.36.0).^70^

To determine TRAP-enriched genes, gene expression was compared between bound and unbound fractions. We used statistical cutoffs for astrocytes (p-value < 0.05, |fold change| > 2) and microglia (p-value < 0.05, |fold change| > 2) to determine input-enriched genes for each cell type for downstream analysis. For analysis of differentially expressed genes (DEGs) between deprived and non-deprived samples, we restricted analysis to the input-enriched genes and used statistical cutoffs p-value < 0.05 and |fold change| > 1.2 for both astrocytes and microglia. Analysis of upstream regulators and root regulators was performed on astrocyte DEGs with the use of QIAGEN IPA (QIAGEN Inc., https://digitalinsights.qiagen.com/IPA).^71^ Ligand-receptor analysis was performed in RStudio using NicheNet (https://github.com/saeyslab/nichenetr).^24^ NicheNet analysis of microglia-astrocyte ligand-receptor interactions was performed using astrocyte DEGs as the “target” gene list, astrocyte input-enriched genes as the “receiver-expressed” gene list and “background” gene list, and microglia input-enriched genes as the “sender-expressed” gene list. To compare the results of this study to publicly available astrocyte TRAP-Seq datasets, NicheNet was performed for each dataset using astrocyte DEGs as the “target” gene list, astrocyte expressed genes as the “receiver-expressed” gene list and “background” gene list, and all ligands in the NicheNet database to reflect an unknown sender. Putative ligands were ranked based on the area under the precision-recall curve (AUPR).

#### Analysis of Wnt signaling by immunofluorescence

Analysis of TCF/Lef:H2B-GFP reporter activity and nuclear β-catenin were performed using single plane 63x confocal images of immunolabeled brain sections containing the barrel region using a Zeiss Observer Spinning Disk Confocal microscope equipped with diode lasers (405nm, 488nm, 594nm, 647nm) and ZEN acquisition software (Zeiss). Data analysis was performed by experimenters blind to the identity of the samples. For each animal, images centered within a barrel in layer IV of the barrel cortex were acquired from 6 to 10 FOV per hemisphere. Using FIJI software, images were background subtracted (rolling ball radius 50 pixels) and manually thresholded using the SOX9 channel to identify astrocyte nuclei or using DAPI to identify all nuclei. The Analyze Particles function (particle size 200 pixels to infinity; ImageJ plugin, NIH) was used to measure the area and mean fluorescence intensity (MFI) in the nucleus (SOX9^+^ and/or DAPI^+^ area) of each cell. For each animal, we quantified the average MFI for all cells in each hemisphere with area > 15 µm^2^.

#### In situ hybridization

Mice were transcardially perfused with PBS and 4% PFA, then brains were extracted, post-fixed in 4% PFA, and cryo-embedded in a 2:1 mixture of OCT : 30% sucrose as described above (see Immunohistochemistry). 10 µm thick coronal sections were cut on a cryostat and mounted on glass slides. *In situ* hybridization was performed using the RNAscope® Multiplex Fluorescent Reagent Kit v2 with TSA Vivid Dyes (323270 Advanced Cell Diagnostics; Newark, CA) following manufacturer’s instructions using probes Mm-Wls-C2 (405011-C2 Advanced Cell Diagnostics) and Mm-P2ry12-C3 (317601-C3 Advanced Cell Diagnostics). On each slide, one section was hybridized with the RNAscope 3-plex Negative Control Probe (320871 Advanced Cell Diagnostics). Briefly, slides were washed with PBS for 5 minutes, baked for 30 minutes at 60°C, post-fixed in 4% PFA for 15 minutes at 4°C, and dehydrated in ethanol. Slides were then incubated in RNAscope hydrogen peroxide for 10 minutes at room temperature, washed in deionized H_2_O, steamed in 99°C RNAscope Target Retrieval solution for 15 minutes, washed in deionized H_2_O and 100% ethanol, and dried at 60°C for 5 minutes. Individual brain sections were outlined on slides using a hydrophobic pen and dried overnight at room temperature. Slides were then treated with RNAscope Protease III for 30 minutes at 40°C, washed with deionized H_2_O, incubated with probes for 2 hours at 40°C, and washed in RNAscope wash buffer. Sections were then hybridized with RNAscope Multiplex FL v2 AMP1, AMP2, and AMP3, labeled with TSA Vivid 570 (1:1500) for C2 and TSA Vivid 650 (1:1500) for C3, and mounted with Fluoroshield mounting media containing DAPI (F6057 Sigma-Aldrich).

Single plane 40x confocal images were acquired using a Zeiss Observer Spinning Disk Confocal microscope equipped with diode lasers (405nm, 488nm, 594nm, 647nm) and ZEN acquisition software (Zeiss). Images were taken of 6-10 FOV per animal from sections labeled with *Wls* and *P2ry12* RNAscope probes and 2 FOV from sections labeled with negative control probes. Data analysis was performed by experimenters blind to the identity of the samples. Using FIJI software, images were background subtracted for each animal such that < 8 dots per FOV were visible in the negative control images. Images were then manually thresholded using *P2ry12* signal to identify microglia. The total *P2ry12^+^*area and *Wls^+^ P2ry12^+^* area were recorded for each image, and the percentage of *P2ry12^+^* area containing *Wls*^+^ puncta was calculated for each animal.

#### XAV939 injections

For inhibition of canonical Wnt signaling, mice were injected once daily i.p. with 7.5 mg/kg XAV939 (X3004 Sigma-Aldrich) dissolved in 10% DMSO at ages P3, P4, P5, P6, P7, P8, and P9. Vehicle-injected mice were injected with 10% DMSO.

#### Intracerebroventricular (icv) injections

To label astrocytes with smV5, P0-P1 mice were anesthetized and injected with two bilateral 2 µL injections of PHP.EB::GfaABC1D-lck-smV5-4x6T adeno-associated virus (AAV) at a titer of 6.37 x 10^13^ viral genomes/mL using a hamilton syringe with a 33 gauge needle. Pups were placed on heat pads for recovery.

### Quantification and Statistical Analysis

Statistical analyses were performed using GraphPad Prism 10.0.3. The exact number of samples (*n*), what *n* represents, the statistical tests used, and *p* values for each experiment are noted in the figure legends. For pairwise comparisons between control and deprived hemispheres, we used the ratio paired t test. For comparisons between two unpaired groups, we used Student’s t-test. For 2-way repeated measures comparisons, we used 2-way repeated measures ANOVA followed by Sidak’s post-hoc test.

---

**Figure S1.**
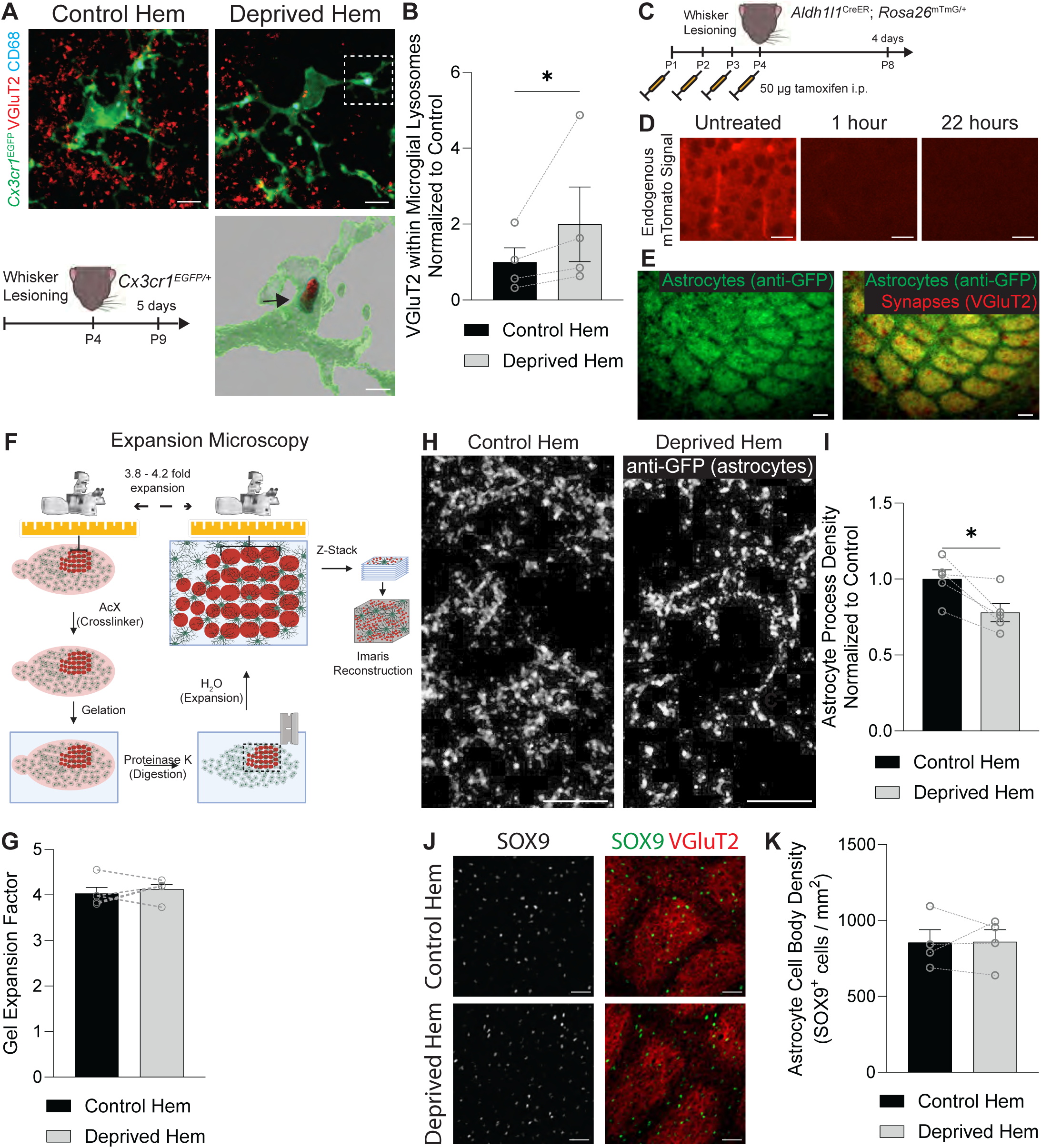
Microglial synapse engulfment is increased and astrocyte process density is reduced after whisker lesioning. Related to Figure 1 (A) Top: representative fluorescent max intensity projection images of anti-VGluT2^+^ thalamocortical synaptic terminals (red), anti-CD68^+^ lysosomes (cyan), and *Cx3cr1^EGFP+^* microglia immunolabeled with anti-GFP (green) in layer IV of the barrel cortex in the control and deprived hemispheres at 5 days post-lesioning. Scale bars 5 µm. Bottom left: diagram of experimental protocol used to assess microglial engulfment of thalamocortical synaptic terminals in *Cx3cr1^EGFP/+^* mice at 5 days after unilateral whisker lesioning. Bottom right: Imaris surface reconstruction of anti-VGluT2^+^ synaptic material (red) contained within anti-CD68^+^ lysosomes (cyan) inside *Cx3cr1^EGFP/+^* microglia immunolabeled with anti-GFP (green) from the white outlined region. Arrow indicates an example of VGluT2^+^ material contained within CD68^+^ microglial lysosomes. Scale bar 2 µm. (B) Quantification of anti-VGluT2^+^ synaptic material within microglial lysosomes at 5 days after whisker lesioning shows more engulfed anti-VGluT2^+^ material in the deprived hemisphere (hem) of the barrel cortex compared to the control hem (Ratio paired t test: *n* = 4 mice. * *p* < 0.05). (C) Diagram of experimental protocol used to assess astrocyte-synapse contacts by expansion microscopy in *Aldh1l1^CreER^; Rosa26^mTmG/+^* mice at 4 days after unilateral whisker lesioning. (D) Representative fluorescent images of endogenous mTomato signal in untreated brain sections from *Aldh1l1^CreER^; Rosa26^mTmG/+^* mice and after treatment with liberate antibody binding (LAB) solution. The endogenous mTomato signal is no longer detectable after 22 hours of incubation in LAB solution. Scale bars 20 µm. (E) Representative fluorescent images of layer IV barrel cortex brain sections from *Aldh1l1^CreER/+^*_;_ *Rosa26^mTmG/+^*mice collected 4 days after unilateral whisker lesioning. Sections from the control and deprived hemispheres are labeled with anti-GFP to identify astrocyte processes and anti-VGluT2 to label thalamocortical synaptic terminals. Scale bars 20 µm. (F) Diagram of experimental protocol to perform expansion microscopy. First, pre-expansion images are taken of brain sections from *Aldh1l1^CreER^; Rosa26^mTmG/+^* mice immunolabeled with anti-GFP to detect astrocytes and anti-VGluT2 to detect thalamortical synaptic terminals in the barrel cortex. Sections are then incubated with the crosslinker Acryloyl-X SE (AcX), embedded in a gel, digested with proteinase K, cut down to the region of interest, and incubated in H_2_O to isotopically expand the gel by ∼3.8 - 4.2 times its original size. A post-expansion image is taken and measurements between specific landmarks are compared with the pre-expansion image to determine the gel expansion factor for each gel. Confocal z-stack images are taken for surface reconstructions using Imaris software. (G) Quantification of the gel expansion factor in the control and deprived hems of *Aldh1l1^CreER^; Rosa26^mTmG/+^* mice shows no differences between hems (Ratio paired t test: *n* = 5 mice). (H) Representative fluorescent images of expanded layer IV barrel cortex brain sections from *Aldh1l1^CreER/+^*_;_ *Rosa26^mTmG/+^* mice collected 4 days after unilateral whisker lesioning. Sections from the control and deprived hem are labeled with anti-GFP to identify astrocyte processes. Scale bars 5 µm (corrected for expansion index). (I) Quantification of astrocyte process density in layer IV of the barrel cortex at 4 days after whisker lesioning shows reduced density in the deprived hem compared to the control hem (Ratio paired t test: *n* = 5 mice. * p < 0.05) (J) Representative fluorescent images from brain sections containing layer IV of the barrel cortex from the deprived and control hem of *Aldh1l1^CreER^; Rosa26^mTmG/+^*mice at 4 days after whisker lesioning. Sections were immunolabeled with anti-SOX9 to label astrocyte nuclei and anti-VGluT2 to label thalamocortical synaptic terminals. Scale bars 50 µm. (K) Quantification of the density of astrocytes (anti-SOX9^+^ cells) in layer IV of the barrel cortex at 4 days after whisker lesioning does not show a difference between the control and deprived hems (Ratio paired t test: *n* = 4 mice). All data are presented as mean ± SEM.

**Figure S2.**
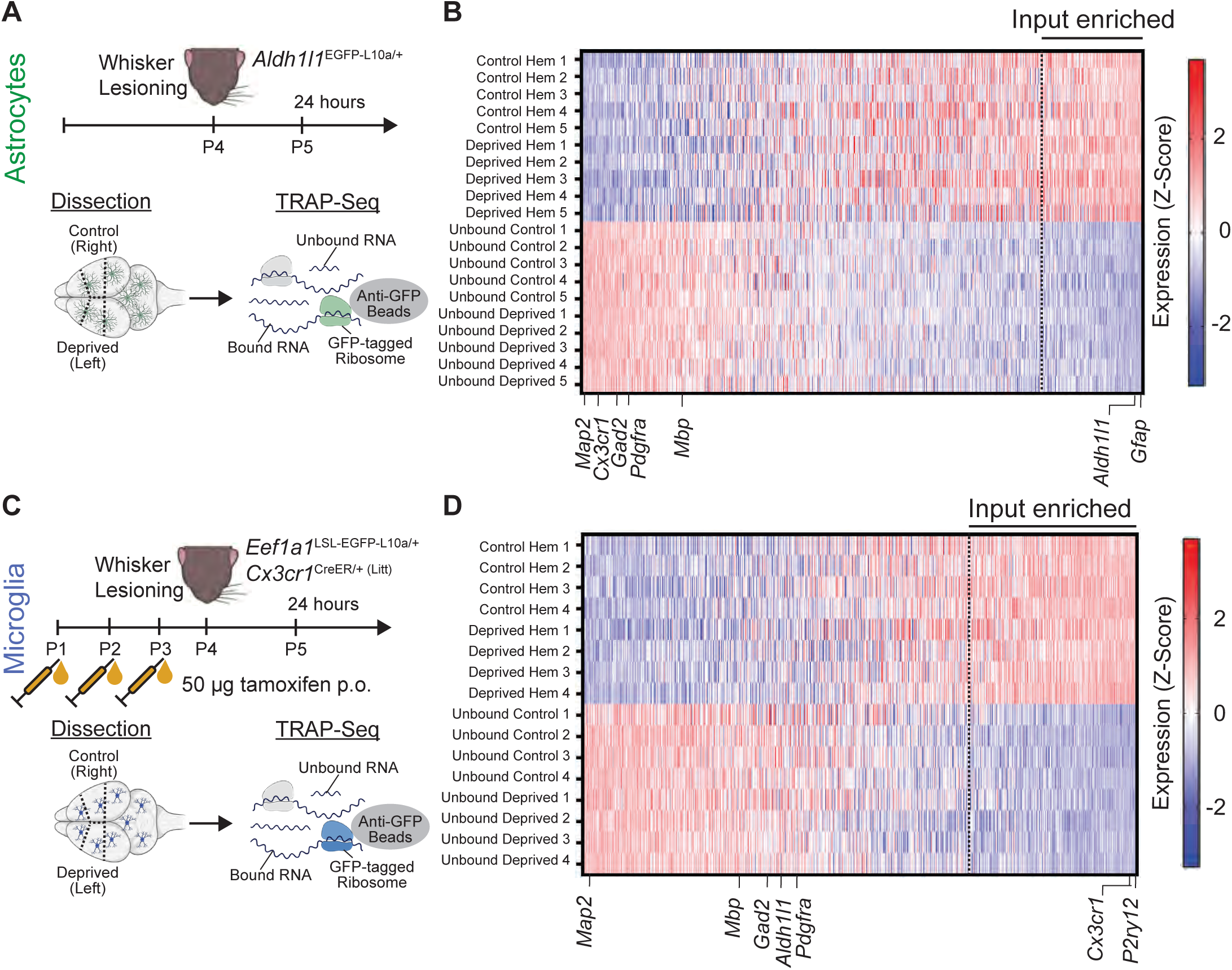
Enrichment of cell-type specific mRNA by TRAP-Seq. Related to Figure 3 (A, C) Diagram of experimental protocol used for TRAP-Seq analysis of ribosome-bound RNA from (A) astrocytes and (C) microglia isolated from the control and deprived hemispheres (hem) of the primary somatosensory cortex at 24 hours after unilateral whisker lesioning. (B, D) Heatmaps showing expression levels of all genes detected in (B) astrocyte and (D) microglia TRAP-Seq experiments, sorted by enrichment in TRAP-enriched samples compared to unbound samples. Rows correspond to individual TRAP-enriched and unbound samples from control and deprived hems. Dashed black lines indicate the cutoffs used to determine input-enriched genes for downstream analyses for astrocytes (p-value < 0.05, |fold change| > 1.2, and mean expression > 5) and microglia (p-value < 0.05, |fold change| > 2, and mean expression > 5). Cell-type specific markers for astrocytes (*Aldh1l1*, *Gfap*), microglia (*Cx3cr1*, *P2ry12*), neurons (*Map2*), interneurons (*Gad2*), oligodendrocytes (*Mbp*), and OPCs (*Pdgfra*) are indicated. Only astrocyte cell-type markers are input-enriched in (B) and only microglia cell-type markers are input-enriched in (D), confirming that TRAP-isolations successfully enriched for the cell type of interest.

**Figure S3.**
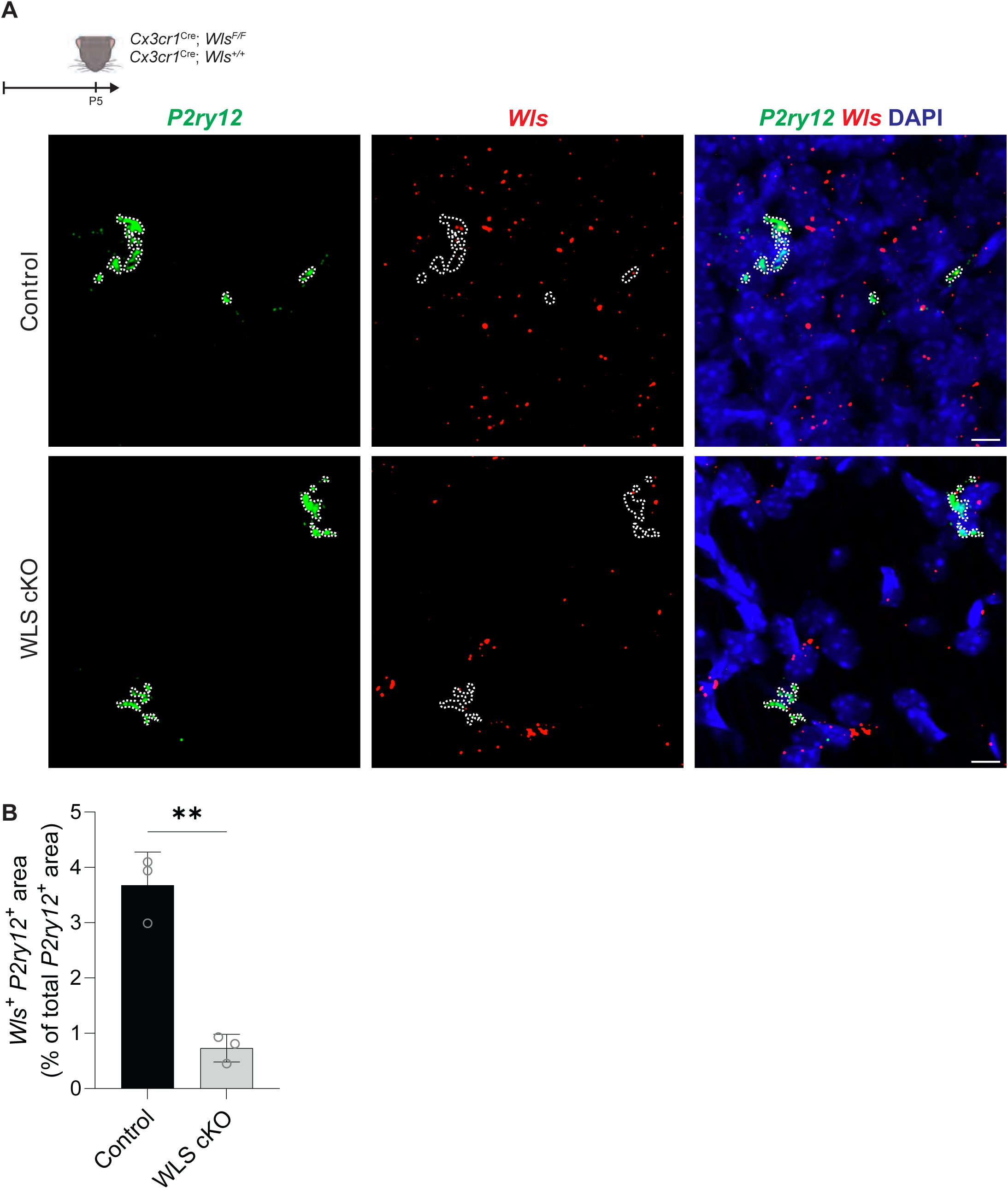
Ablation of *Wls* in WLS cKO mice. Related to Figure 5 (A) (Top) diagram of experimental protocol used to perform *in situ* hybridization of *Wls* in *Cx3cr1^Cre^; Wls^F/F^* (WLS cKO) and *Cx3cr1^Cre^; Wls^F/F^* (control) mice at postnatal day 5 (P5). (Bottom) representative fluorescent images of brain sections from WLS cKO and control mice with *in situ* hybridization of *Wls* (red) and the microglia marker *P2ry12* (green). Nuclei are labeled with DAPI (blue). White outlines indicate the *P2ry12^+^* area. Scale bars 10 µm. (B) Quantification of the percentage of *P2ry12^+^* area that overlaps with *Wls* fluorescence in WLS cKO mice and control mice at P5 shows reduced *Wls*^+^ area in WLS conditional mice, confirming that *Wls* was successfully ablated (Student’s t test: *n* = 3 WLS cKO mice, 3 control mice. ** *p* < 0.01). All data are presented as mean ± SEM.

**Figure S4.**
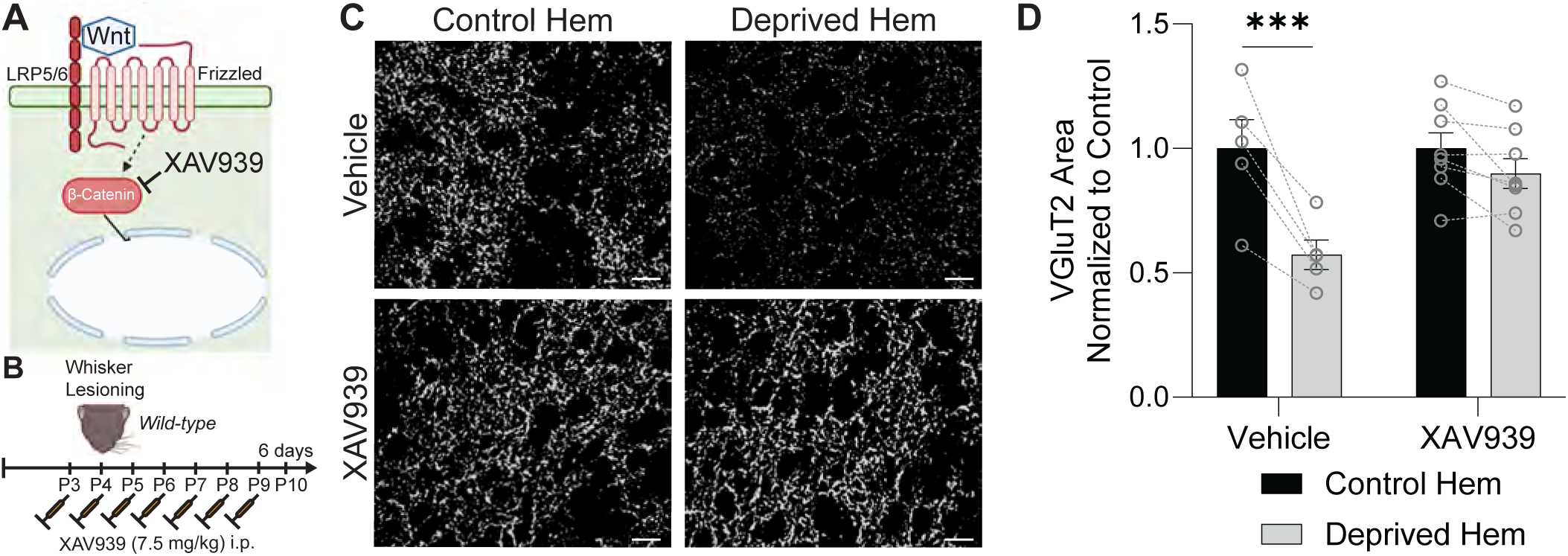
A pharmacological inhibitor of Wnt receptor signaling prevents synapse loss after whisker lesioning. Related to Figure 5 (A) Diagram of the effect of XAV939 on the canonical Wnt siganling pathway. XAV939 promotes the degradation of β-catenin, which prevents it from accumulating in the nucleus after binding of Wnt ligand to frizzled and LRP5/6 receptors. (B) Diagram of experimental protocol used to assess the density of VGluT2^+^ synaptic terminals in layer IV of the barrel cortex at 6 days after whisker lesioning in wild-type mice injected with the Wnt signaling inhibitor XAV939. (C) Representative fluorescent images of anti-VGluT2^+^ thalamocortical synaptic terminals in layer IV of the barrel cortex in the control and deprived hemispheres (hem) at 6 days post-lesioning in wild-type mice injected with XAV939 or vehicle. Scale bars 10 µm. (D) Quantification of the density of anti-VGluT2^+^ thalamocortical synaptic terminals in layer IV of the barrel cortex at 6 days post-lesioning shows reduced density in the deprived hem compared to the control hem in vehicle treated mice, but not in mice treated with XAV939 (Repeated measures 2-way ANOVA with Sidak’s post hoc test: *n* = 5 vehicle mice, 8 XAV939 mice. *** *p* < 0.001). All data are presented as mean ± SEM.

**Figure S5.**
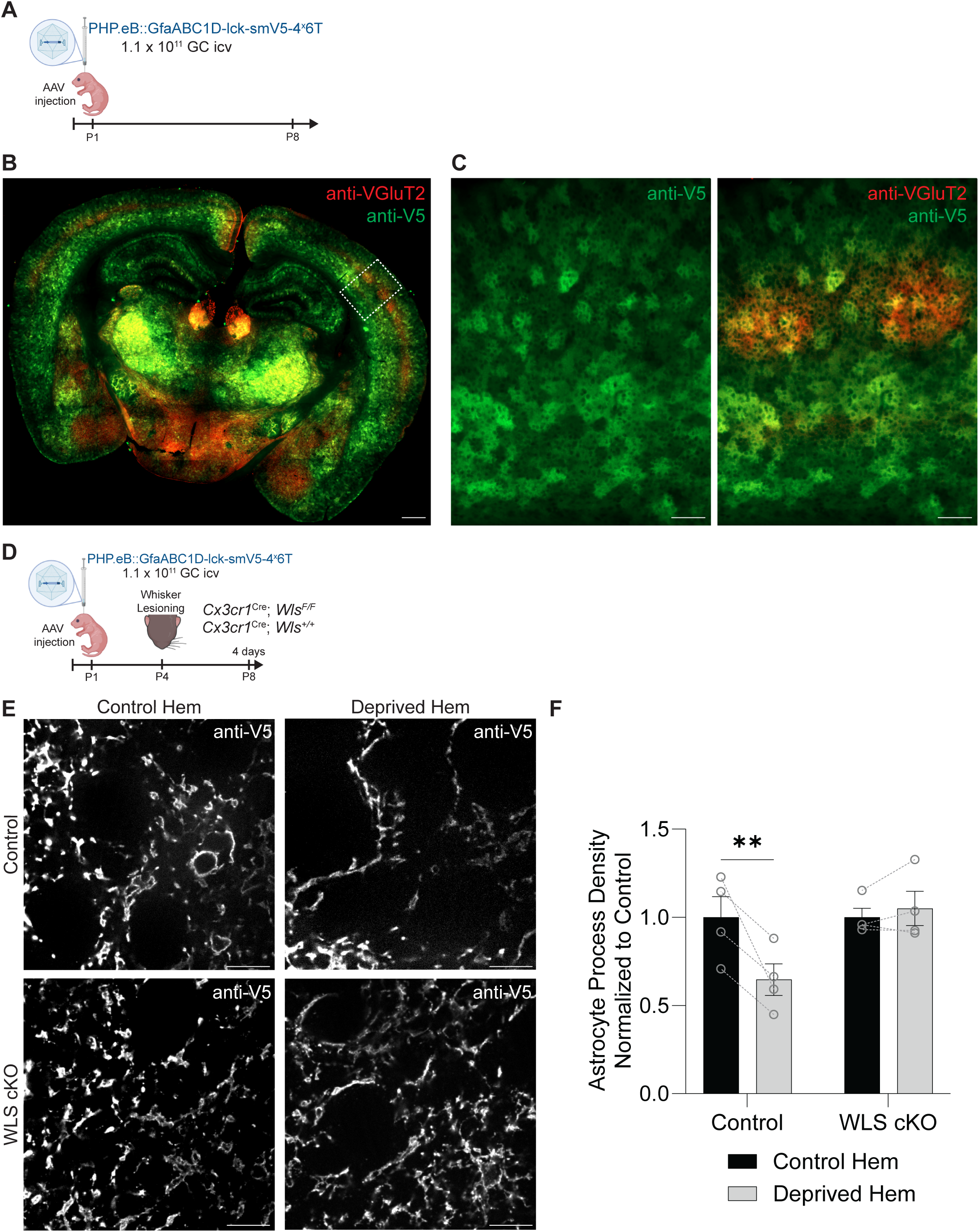
Ablation of microglial Wnt release blocks reductions in astrocyte process density after whisker lesioning. Related to Figure 5 (A) Diagram of experimental protocol used to label astrocytes with membrane-bound V5 protein at postnatal day 8 (P8) by intracerebroventricular (icv) injection of PHP.EB::GfaABC1D-lck-smV5-4^x^6T adeno-associated virus (AAV). (B-C) Representative fluorescent images of brain sections from P8 mice injected with PHP.EB::GfaABC1D-lck-smV5-4^x^6T AAV, immunolabeled with anti-VGluT2 (red) and anti-V5 (green). White box region in (B) is shown at higher magnification in (C). Scale bars (B) 500 µm and (C) 100 µm. (D) Diagram of experimental protocol used to assess astrocyte-synapse interactions in *Cx3cr1^Cre^; Wls^F/F^* (WLS cKO) and *Cx3cr1^Cre^; Wls^F/F^* (control) mice at 4 days after unilateral whisker lesioning. Astrocytes were labeled with membrane-bound V5 protein by icv injection of PHP.EB::GfaABC1D-lck-smV5-4^x^6T AAV at P1. (E) Representative fluorescent images of expanded layer IV barrel cortex brain sections from WLS cKO and control mice collected 4 days after unilateral whisker lesioning. Sections from the control and deprived hemispheres are labeled with anti-V5 to identify astrocyte processes. Scale bars 5 µm (corrected for expansion index). (F) Quantification of astrocyte process density in layer IV of the barrel cortex at 4 days after unilateral whisker lesioning shows reduced process density in the deprived hem of control mice compared to the control hem, but not in WLS cKO mice (Repeated measures 2-way ANOVA with Sidak’s post-hoc test: *n* = 4 control mice, 4 WLS cKO mice. ** *p* < 0.01). All data are presented as mean ± SEM.

